# Characterizing the rates and patterns of *de novo* germline mutations in the aye-aye (*Daubentonia madagascariensis*)

**DOI:** 10.1101/2024.11.08.622690

**Authors:** Cyril J. Versoza, Erin E. Ehmke, Jeffrey D. Jensen, Susanne P. Pfeifer

## Abstract

Given the many levels of biological variation in mutation rates observed to date in primates – spanning from species to individuals to genomic regions – future steps in our understanding of mutation rate evolution will be aided by both a greater breadth of species coverage across the primate clade, but also by a greater depth as afforded by an evaluation of multiple trios within individual species. In order to help bridge these gaps, we here present an analysis of a species representing one of the most basal splits on the primate tree (aye-ayes), combining whole-genome sequencing of seven parent-offspring trios from a three-generation pedigree with a novel computational pipeline that takes advantage of recently developed pan-genome graphs, thereby circumventing the application of (highly subjective) quality metrics that has previously been shown to result in notable differences in the detection of *de novo* mutations, and ultimately estimates of mutation rates. This deep sampling has enabled both a detailed picture of parental age effects as well as sex dependency in mutation rates which we here compare with previously studied primates, but has also provided unique insights into the nature of genetic variation in one of the most endangered primates on the planet.

## INTRODUCTION

As the ultimate source of novel genetic variation, a comprehensive understanding of mutational processes is a requisite for interpreting rates and patterns of molecular evolution. Partly for anthropocentric reasons, considerable attention has been paid to studying the causes and consequences of mutations in humans specifically, not least to improve the dating of events in our species’ evolutionary history (Nielsen et al. 2017), infer phylogenetic relationships with other closely-related primates (Kuderna et al. 2023), and improve our understanding of the impact of the mutational process on health and disease (Shendure and Akey 2015).

Prior to the genomic age, the inference of mutation rates relied on indirect observations; for example, by estimating rates based on the frequency of newly arising autosomal dominant or X-linked recessive Mendelian diseases (Haldane 1935, 1947; Kondrashov 2003; Nachman 2004; Lynch 2010). With early genetic data, neutral sequence divergence between two closely-related species could additionally be utilized (e.g., humans and chimpanzees; Nachman and Crowell 2000; Chimpanzee Sequencing and Analysis Consortium 2005) – a strategy that relies on the ’clock-like’ accumulation of these fixations due to the fact that the rate of neutral divergence is dictated by the rate of neutral mutation (Kimura 1968, 1983). Despite providing highly useful insights, estimates obtained from both approaches are also fraught with substantial uncertainty (see review by Drake et al. 1998), given that the resulting parameter estimates can be compromised if the underlying assumptions are violated (e.g., if the mutational target size of the Mendelian disease in question is large, or if phylogenetic calibration rates – required to convert substitutions accumulated between lineages to divergence times – have not remained constant throughout evolutionary history, respectively).

However, progress in sequencing technologies and computational methodologies has newly enabled researchers to investigate genomes at scale. It is thus now feasible to characterize the rates and patterns of contemporary spontaneous (*de novo*) germline mutation (DNM) in a direct and relatively comprehensive manner, by comparing the genetic code of parents and their offspring (i.e., parent-offspring trios; see review of Pfeifer 2020). As a result, the past years have witnessed notable advances in our understanding of DNMs in humans and non-human primates (Roach et al. 2010; Conrad et al. 2011; Campbell et al. 2012; Kong et al. 2012; Michaelson et al. 2012; Venn et al. 2014; Francioli et al. 2015; Besenbacher et al. 2016; Goldmann et al. 2016; Rahbari et al. 2016; Wong et al. 2016; Jónsson et al. 2017; Pfeifer 2017a; Tatsumoto et al. 2017; Thomas et al. 2018; Besenbacher et al. 2019; Sasani et al. 2019; Kessler et al. 2020; Wang et al. 2020; Wu et al. 2020; Bergeron et al. 2021; Campbell et al. 2021; Yang et al. 2021) as well as in a multitude of other model and non-model organisms (e.g., Bergeron et al. 2023). These studies have highlighted substantial variation in rates between species – including several fold among primates (see reviews by Tran and Pfeifer 2018; Chintalapati and Moorjani 2020) and by orders of magnitude across the Tree of Life (see reviews by Baer et al. 2007; Pfeifer 2020) – yet, our understanding of the biological mechanisms facilitating this evolution across taxa still remains limited.

Germline point (i.e., single nucleotide) mutations are thought to predominantly originate from copying errors during DNA replication left uncorrected by cellular repair mechanisms during spermatogenesis and early embryonic development (see review of Beichman et al. 2024). Due to the nature of gametogenesis, sex-specific differences in the accumulation of replication-driven germline mutations are thus to be expected from first principles (Crow 2000). Specifically, corresponding with a larger number of germline cell divisions in males compared to females, a male mutation bias – that is, a greater contribution of DNMs originating from the paternal compared to the maternal germline (Haldane 1935, 1947; Crow 2000, 2006) – has been observed in many species (Ellegren 2007; Wilson Sayres et al. 2011). Additionally, as spermatogenesis continues throughout adulthood, evidence suggests that this male mutational burden increases with paternal age (*i.e.*, paternal age effect; see reviews by Ségurel et al. 2014; Goriely 2016). However, recent research has demonstrated that there is also a much less-pronounced maternal age effect, suggesting that spontaneous, replication-independent DNA damage in gametes – caused, for example, by extrinsic mutational agents arising from environmental exposure to chemical mutagens and ultraviolet radiation – also plays an important role in the genesis of mutations (Goldmann et al. 2016; Wong et al. 2016; Gao et al. 2019; Wu et al. 2020).

Further contributing to differences in mutation rates are the biochemical mechanisms underlying DNA replication fidelity and repair efficiency (Driscoll and Migeon 1990; and see review by Mohrenweiser et al. 2003). Given that these processes can vary considerably based on genomic features, chromatin state, and the timing of replication, they play a critical role in determining the mutation rates along different regions of the genome (Tyekucheva et al. 2008). Most noteworthy in this regard, CpG sites have an order of magnitude higher *de novo* mutation rate than non-CpG sites in primates, owing to spontaneous methylation-dependent deamination that leads to higher rates of C-to-T transitions (Nachman and Crowell 2000; Hwang and Green 2004; Leffler, Gao, Pfeifer, Ségurel et al. 2013; and see review by Hodgkinson and Eyre-Walker 2011). As a result, whereas the vast majority of germline mutations accrue at a rate proportional to the generation time, inefficiently repaired replication-independent CpG transitions appear to instead accumulate in a more clock-like manner proportional to absolute time (Gao et al. 2016; Moorjani et al. 2016a). Notably however, a paternal age effect has also been observed for CpG>TpG mutations in humans, suggesting that there is no strict molecular clock (Jónsson et al. 2017).

Given the considerable biological variation of mutation rates observed at these multiple scales in primates – spanning from species to individuals to genomic regions (see review by Ségurel et al. 2014) – it will thus be highly informative to expand upon earlier work both by sampling broadly across the primate clade (i.e., outside of the great apes and biomedically-relevant species such as vervet monkeys (Pfeifer 2017a), owl monkeys (Thomas et al. 2018), rhesus macaques (Wang et al. 2020; Bergeron et al. 2021), and baboons (Wu et al. 2020)), as well as by evaluating multiple trios within individual species (i.e., single trios remain the norm, leaving individual-level variation largely unexamined; Bergeron et al. 2023). Such studies will be essential not only for quantifying the degree of mutation rate evolution over deep time-scales, but also for evaluating hypotheses pertaining to the forces governing such change (Sung et al. 2012; Lynch et al. 2016; and see review of Beichman et al. 2024).

One species of particular interest in this comparative regard is the aye-aye (*Daubentonia madagascariensis*) – one of the most basal extant primates. The aye-aye, a strepsirrhine primate that inhabits dry, deciduous, and rain forests of Madagascar – also stands on the verge of extinction (Randimbiharinirina et al. 2019), thus rendering studies of variation of great practical interest at the conservation level as well. As a solitary species that requires extensive individual home territories (often in excess of 1,000 hectares), aye-aye populations are severely threatened by the continued anthropogenic destruction of their habitats. In particular, deforestation from slash-and-burn agriculture, illegal logging, and mining, which have jointly led to the loss of more than 80% of the island’s natural biotope over the past decades (Suzzi-Simmons 2023), are thought to have coincided with a massive decline (≥ 50%) in wildlife populations (Louis et al. 2020). As a result, aye-ayes are now classified as one of the 25 most endangered primates in the world (Randimbiharinirina et al. 2019), and the protection of the last individuals remaining in the wild (estimated to be on the order of a few thousand individuals; Mittermeier et al. 2010) is a key priority of contemporary conservation measures in Africa.

One important aspect of such conservation strategies will necessarily involve developing an improved understanding of the evolutionary forces dictating the generation and maintenance of genetic variation in aye-ayes in the face of small and likely declining population sizes, as this variation will partly dictate the future success of this species. Population genomics allows for the investigation and quantification of these forces dictating levels of variation in this species, providing insights into rates and patterns of mutation as discussed here, structural variation (Versoza et al. 2024), recombination (Versoza, Lloret-Villas et al. 2024), genetic drift as dictated by population history (Terbot et al. 2024), and natural selection (Soni et al. 2024). Importantly, combining this evolutionary genomic information with ecological surveys and behavioral data can be utilized to facilitate on-going efforts to conserve both self-sustaining wild populations as well as populations in captivity.

By combining deep whole-genome sequencing of seven parent-offspring trios from a three-generation pedigree (Figure 1) with a novel computational pipeline that takes advantage of recently developed pan-genome graphs, we thus here characterize the rates and patterns of *de novo* germline mutations in the aye-aye. In addition, the long reproductive life span of aye-ayes – ranging from sexual maturity at 8-36 months of age (Winn 1994) to more than 30 years (Zehr et al. 2014) – allows us to obtain a detailed picture of parental age effects and sex dependency in this highly endangered species, and to compare patterns with those previously observed in other primates.

**Figure 1.**
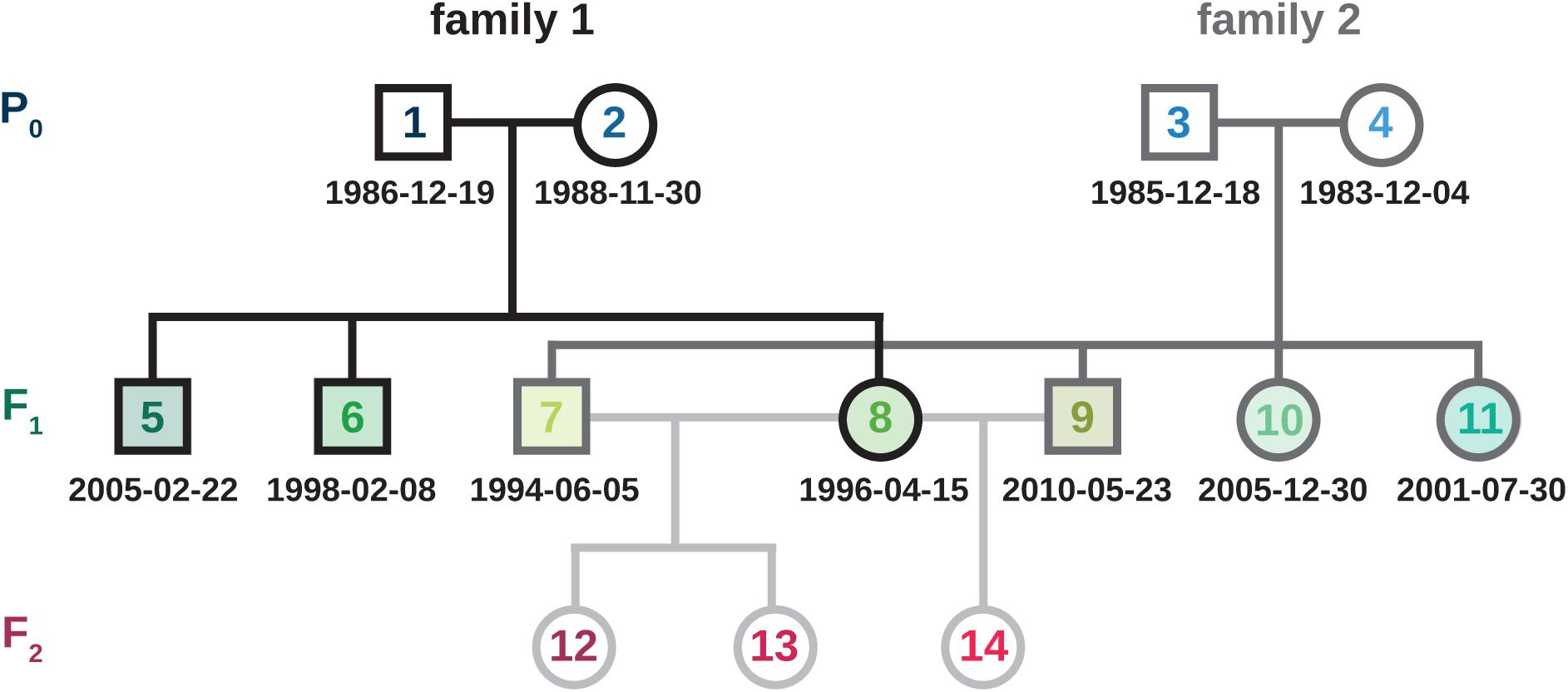
Structure of the aye-aye pedigree. The pedigree was comprised of a parental (P_0_) generation (shown in blue), consisting of two sires and two dams that had a total of seven focal (F_1_) offspring (three and four offspring per breeding pair in families 1 and 2, respectively; shown in green) as well as three offspring of a third (F_2_) generation (shown in red). Squares and circles represent males and females, respectively. The date of birth (yyyy-mm-dd) of the parental and focal individuals is provided underneath the symbols.

## RESULTS AND DISCUSSION

### Identification of DNMs in parent-offspring trios

The genomes of 14 aye-ayes (*D. madagascariensis*) from a three-generation pedigree were sequenced to an average coverage of 52X (range: 48.5X to 54.5X; Supplementary Table 1). The pedigree was comprised of a parental (P_0_) generation, consisting of two sires and two dams that had a total of seven focal (F_1_) offspring (three and four offspring per breeding pair in families 1 and 2, respectively) which were used to identify DNMs in the parent-offspring trios (Figure 1). The age of the P_0_ individuals at the time of birth of their offspring ranged from 7.4 to 26.5 years in females and from 8.5 to 24.4 years in males, spanning the majority of the reproductive life span of the species (Winn 1994; Zehr et al. 2014). Additionally, inclusion of a third (F_2_) generation, composed of three offspring of three of the F_1_ individuals, enabled the investigation of DNM transmission to the next generation. This information also aids in the distinction between mutations that occurred in the germline from those that originated in the soma (Ségurel et al. 2014).

Following the Genome Analysis Toolkit (GATK) Best Practices for non-model organisms (van der Auwera et al. 2013; van der Auwera and O’Connor 2020), variants were called based on the quality-controlled sequencing reads of each individual mapped to the species-specific genome assembly (Versoza and Pfeifer 2024), and jointly genotyped across samples to improve performance. This variant dataset, consisting of 3.6 million autosomal, biallelic single nucleotide polymorphisms (SNPs) with a transition-transversion ratio (Ts/Tv) of 2.47 across the pedigree (Supplementary Table 2), was limited to 7,907 Mendelian violations observed in the seven trios – that is, sites at which individuals of the P_0_ generation were homozygous for the reference allele while at least one of their focal F_1_ offspring was heterozygous.

As errors in sequencing, mapping, variant calling, and genotyping occur at an order of magnitude greater rate than genuine DNMs in primates (Pfeifer 2021), studies frequently apply stringent quality filtering – based, for example, on read coverage, allelic balance (i.e., the ratio of reads carrying the alternative vs reference alleles), and genome complexity (e.g., excluding highly repetitive regions which are notoriously challenging for read mapping and variant calling) – to weed out false positives from an initial set of Mendelian violations (see review by Beal et al. 2012). However, not only is the selection of such quality metrics highly subjective, it can also result in notable differences in the number of DNMs detected, ultimately resulting in substantial differences in estimated mutations rates (see Bergeron et al. 2022). Moreover, the application of sequence metrics also makes it difficult to obtain an accurate and unbiased estimate of the number of sites accessible to the study (often referred to as "accessible sites" or "callable genome"), necessary to calculate the per-site mutation rate; this is a particular challenge for those filter criteria that do not have an equivalent between variant and invariant sites (for an in-depth discussion, see Pfeifer 2021). To avoid an arbitrary selection of filter criteria, Mendelian violations were instead re-genotyped using a highly accurate pan-genome approach to increase specificity (Eggertsson et al. 2017), resulting in 459 DNM candidates with high-confidence calls of the mutant allele across the seven parent-offspring trios. As validation experiments designed to assess the accuracy of DNMs via orthogonal technologies (such as Sanger sequencing) are challenging in non-model organisms – both in terms of their failure rates (e.g., a similar study in chimpanzees reported an assay failure rate of >20%; Venn et al. 2014) as well as additional sample requirements (which can be problematic, particularly for endangered species) – candidate sites were instead visually inspected for common signs of sequencing, read mapping, variant calling, and/or genotyping errors to guard against mis-genotyping. A total of 323 out of 459 candidate sites passed this manual curation performed independently by two researchers (Supplementary Table 3), indicating a false discovery rate of 29.6%.

Several lines of evidence suggest that these visually validated DNMs are genuine rather than sequencing artifacts. Firstly, as expected for genuine DNMs, none of the 323 validated mutations were found to be segregating in a previous sample of 12 unrelated aye-aye individuals (Perry et al. 2013). Secondly, none of the DNMs were located within regions of structural variation (Versoza et al. 2024) that might have complicated read mapping, potentially leading to spurious variant calls (Sedlazeck et al. 2018). Finally, the transmission rates of DNMs identified in the F_1_ individuals to their F_2_ offspring were consistent with Mendelian expectations (transmission rates ranged from 0.36 to 0.56, with an average of 0.48; Fisher’s exact test: *p*-value = 0.5637), with work by Wang and Zhu (2014) demonstrating that DNMs detected using a three-generation pedigree approach are indeed in agreement with those validated by an orthogonal technology.

### Genomic distribution of DNMs and the mutation spectrum

The identified DNMs were distributed equally across the autosomes, with the majority harbored within intergenic and intronic regions (47.4% and 35.6%; Supplementary Figure 1) as expected from the overall genome composition (χ^2^ = 1.5028, df = 7, *p*-value = 0.9822). In addition, in agreement with the repeat content of the aye-aye genome (35.0%), 106 out of the 323 DNMs (32.8%) were located within repetitive regions (Fisher’s exact test: *p*-value = 0.618). In many organisms including primates, repetitive elements tend to be methylated to maintain genomic integrity (see review by Slotkin and Martienssen 2007), thus leading to frequent C>G transitions at CpG sites in these regions. In fact, higher C>T mutation rates at methylated CpG sites (Hwang and Green 2004; Hodgkinson and Eyre-Walker 2011) account for ∼17-19% of DNMs observed in humans (Kong et al. 2012; Ségurel et al. 2014). Although several non-human primates exhibit lower or higher fractions of CpG>TpG DNMs (e.g., 12% in owl monkeys [Thomas et al. 2018] and 24–29% in chimpanzees [Venn et al. 2014; Tatsumoto et al. 2017]), aye-ayes display a near-identical trend to that observed in humans (17.6%; binomial test: *p*-value = 0.5709). Indeed, after accounting for species-specific differences in CpG>TpG transitions, the mutation spectra of haplorrhines – as assessed from catarrhines (humans [Kong et al. 2012; Besenbacher et al. 2016; Goldmann et al. 2016; Rahbari et al. 2016], chimpanzees [Venn et al. 2014; Besenbacher et al. 2019], gorillas [Besenbacher et al. 2019], orangutans [Besenbacher et al. 2019], vervet monkeys [Pfeifer 2017a], rhesus macaques [Wang et al. 2020], and baboons [Wu et al. 2020]) and platyrrhines (owl monkeys [Thomas et al. 2018]) – is remarkably similar to that of strepsirrhines (as assessed from aye-ayes; χ^2^ = 6.9131, df = 4, *p*-value = 0.1406; Figure 2), suggesting a conservation of the underlying molecular machinery over long evolutionary time scales.

**Figure 2.**
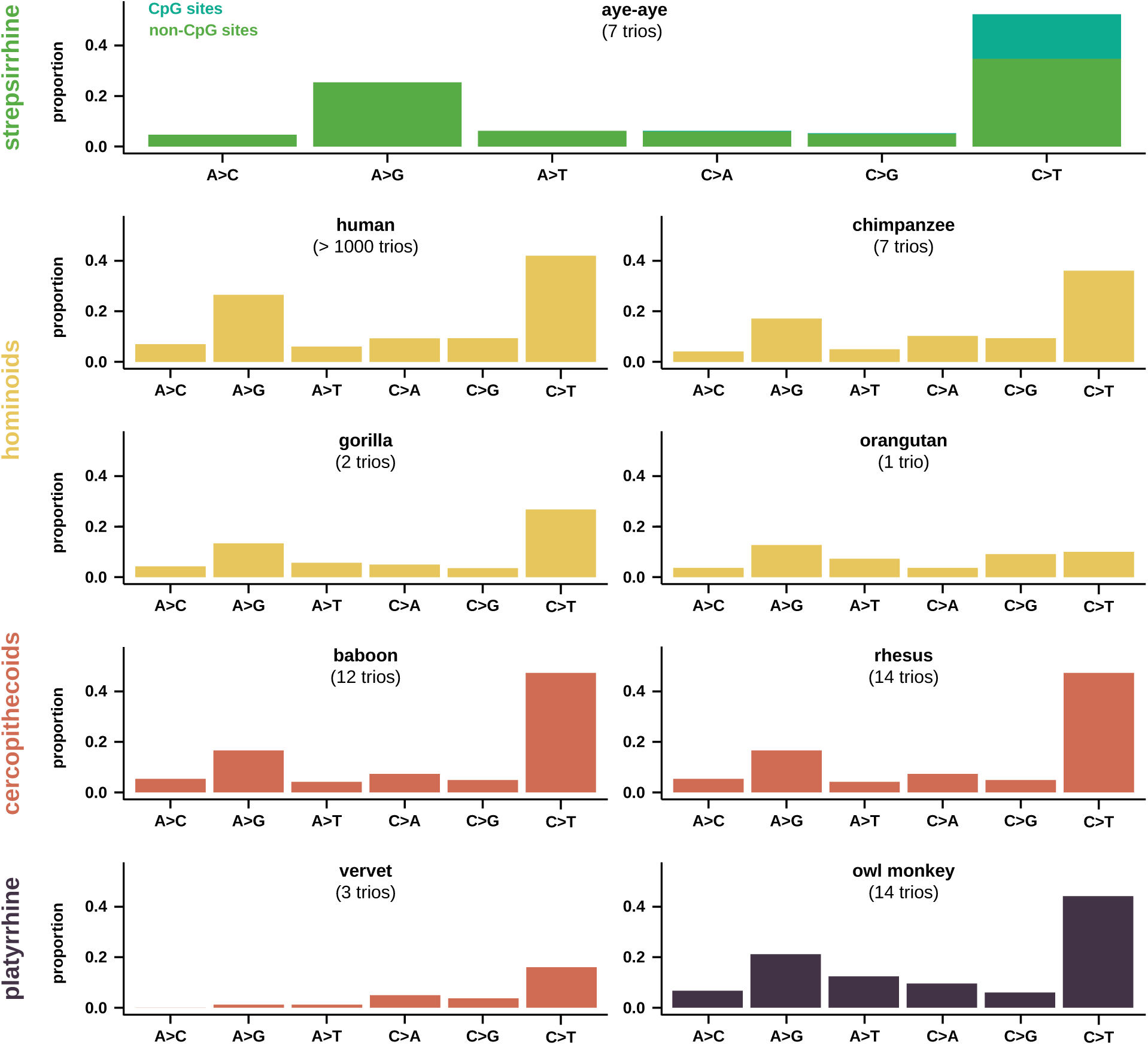
Primate mutation spectra. A comparison of mutation spectra obtained from previously published hominoid (human [Kong et al. 2012; Besenbacher et al. 2016; Goldmann et al. 2016; Rahbari et al. 2016], chimpanzee [Venn et al. 2014; Besenbacher et al. 2019], gorilla [Besenbacher et al. 2019], and orangutan [Besenbacher et al. 2019]; shown in yellow), cercopithecoid (baboon [Wu et al. 2020], rhesus macaque [Wang et al. 2020], and vervet monkey [Pfeifer 2017a]; shown in red), and platyrrhine (owl monkey [Thomas et al. 2018]; shown in purple) datasets for which more than a single trio was available (with the exception of orangutan) as well as aye-ayes (this study) as a representative of the more distantly related strepsirrhines (with mutations at CpG sites shown in teal and at non-CpG sites in green). The relative proportion of each mutation type is shown, with reverse complements collapsed.

### DNM clustering and sibling-sharing

A non-random clustering of DNMs has previously been observed in several primates (Campbell et al. 2012; Michaelson et al. 2012; Venn et al. 2014; Francioli et al. 2015); for example, in a similarly sized chimpanzee pedigree, 17% of DNMs were clustered within 1Mb of another DNM in the six trios studied (Venn et al. 2014). Similarly, in aye-ayes, 10.2% of all DNMs were located within 1Mb of another event, with 27.3% of clustered DNMs originating in the same individual (3 and 2 DNMs within 1kb and 10kb in individual 9, respectively; 2 DNMs within 100kb in individual 10; 2 DNMs within 1Mb in individual 5). These intra-individual clusters of DNMs at nearby locations might potentially result from an error-prone polymerase, inefficient DNA repair, or from a shared exposure to mutagenic agents. Additionally, two DNMs were carried by more than one F_1_ offspring in a family, suggesting that they have likely arisen through mutations that occurred either prior to primordial germ cell specification or during the early post-zygotic stages which are known to be particularly error-prone (Woodruff and Thompson 1992; and see reviews by Biesecker and Spinner 2013; Ségurel et al. 2014; Samuels and Friedman 2015). In agreement with previous observations in humans, the DNMs shared between siblings are CpG>TpG transitions and occurred in the same sibling pair as might be expected given that the sharing of a first DNM has been shown to increase the probability of a subsequent sharing by more than 20% (Jónsson et al. 2018). This observation of sibling-sharing of DNMs reaffirms the importance of mutational processes occurring prior to the final meiotic germ cell division in shaping the mutational landscape (for a detailed discussion, see Scally 2016).

### Estimating germline mutation rates and parental age effects

Translating the number of DNMs to an estimated per-site germline mutation rate requires not only a careful assessment of both the false discovery rate as well as the length of the genome accessible to the study as discussed above, but also a robust estimation of the false negative rate. In order to assess the number of genuine DNMs that might have been missed, 1,000 DNMs were simulated within sequencing reads in a manner that mimicked empirical haplotype structure, read coverage, and allele balance. These simulations were subsequently run through our computational pipeline, yielding a false negative rate of 9.5%. Correcting for the estimated false discovery and false negative rates, inferred per-site germline mutation rates ranged from 0.4 x 10^-8^ in an individual born to young parents (maternal and paternal ages at birth were 9.2 and 11.2 years, respectively) to 2.0 x 10^-8^ in an individual born to old parents (26.5 and 24.4 years), with an average rate of 1.1 x 10^-8^ across the trios studied.

Thus, in aye-ayes, there is strong evidence for a parental age effect on the rate of mutation (paternal age: *r^2^* = 0.7061, *p*-value = 0.0179; maternal age: *r^2^* = 0.8748, *p*-value = 0.0020; Figure 3a). The overall effect was consistent and independent of genomic background; however, the observation was statistically significant only outside of repeats (repeat background: *r^2^* = 0.3857, *p*-value = 0.1367; non-repeat background: *r^2^* = 0.7811, *p*-value = 0.0083; Figure 3b). As maternal and paternal ages at birth are highly correlated in this study (*r* = 0.9603, *p*-value = 0.0006), the parent-of-origin of the DNMs was determined using a combination of direct (read-based) and indirect (genetic) phasing. Although read tracing enabled the phasing of 95.1% of heterozygous variants in the focal offspring on average (range: 93.5– 98.1%), only ∼10% of reads could be resolved by haplotype due to the low levels of genetic diversity in aye-ayes (Perry et al. 2013). Moreover, no heterozygous sites for which the parent-of-origin could unequivocally be determined were located within the paired-end sequencing reads harboring DNMs, and thus read-based phasing was unable to resolve the parental origin of any DNMs inherited by the F_1_ individuals. This finding was anticipated given that even in primate species with much higher levels of nucleotide diversity (such as chimpanzees) only a small fraction of DNMs (∼25%) can generally be phased using short-read data (Venn et al. 2014). However, genetic phasing based on the transmission of haplotypes across the three-generation pedigree could be used to assign the parental origin of DNMs carried by the two F_1_ individuals with multiple offspring (i.e., individuals 7 and 8). Out of the 37 phased DNMs, 27.0% and 73.0% were found to be maternal and paternal in origin, respectively. Moreover, despite the dataset being small, more C>T DNMs of maternal than paternal origin were observed (50.0% vs 40.7%), consistent with earlier observations in humans (Goldmann et al. 2016; Jónsson et al. 2017).

**Figure 3.**
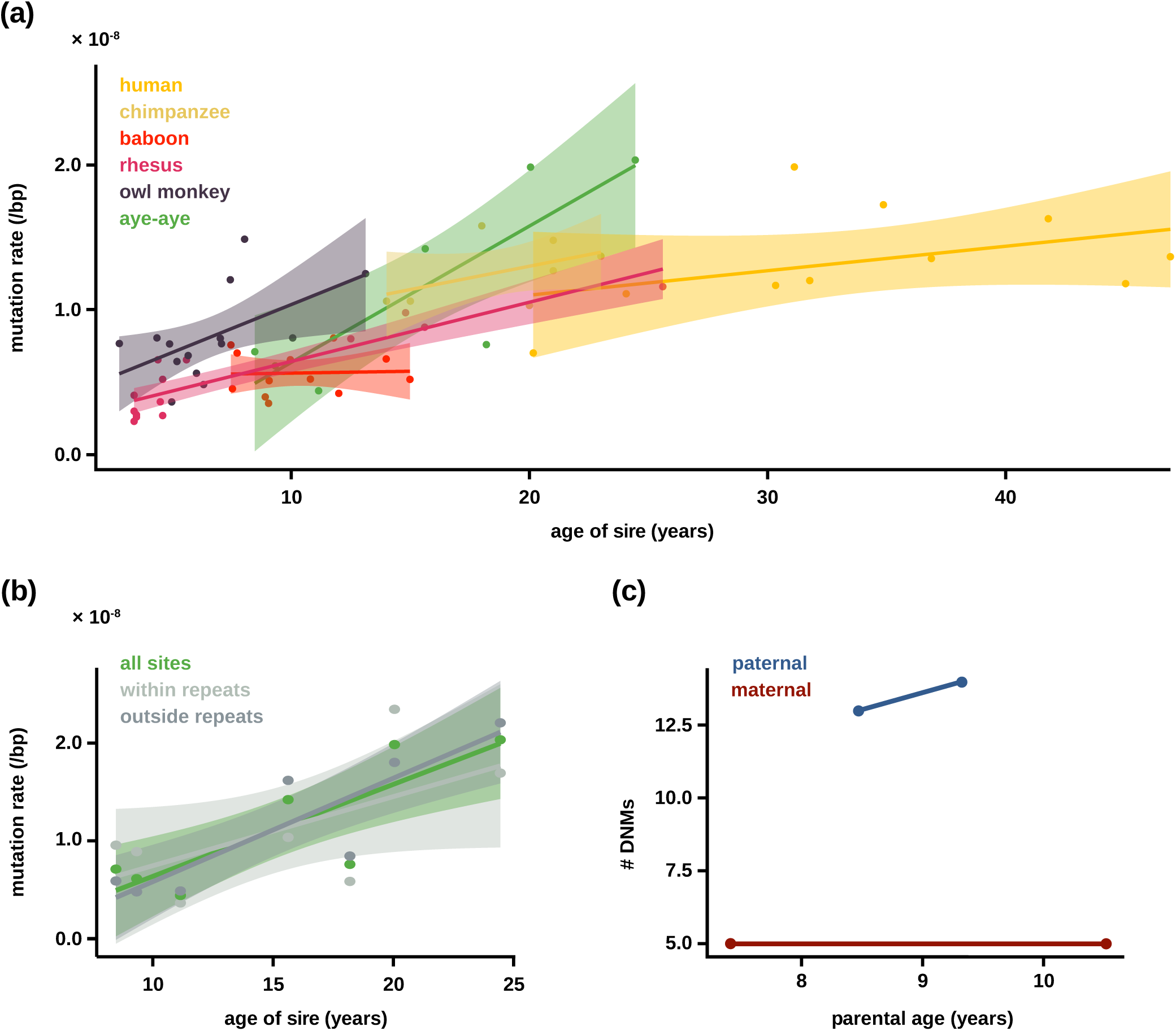
Primate mutation rate estimates. Mutation rate estimates obtained from previously published haplorrhines (including both catarrhines and platyrrhines), compared to aye-ayes as a representative of strepsirrhines. (a) Relationship between paternal age at birth (in years) and mutation rate (per base pair [bp]) in humans (shown in yellow; Wu et al. 2020), chimpanzees (gold; Venn et al. 2014; Besenbacher et al. 2019), baboons (red; Wu et al. 2020), rhesus macaques (pink; Wang et al. 2020), owl monkeys (purple; Thomas et al. 2018), and aye-ayes (green; this study). Linear regression and 95% confidence intervals are shown as solid lines and shaded areas, respectively. (b) Relationship between paternal age at birth and mutation rates in aye-ayes at all sites (shown in green) as well as within and outside of repetitive genomic regions (light and dark gray, respectively). (c) Relationship between parental age at birth and the number of phased *de novo* mutations (# DNMs) of maternal and paternal origin (shown in red and blue, respectively).

The male mutation bias of 2.6-2.8 observed in aye-ayes is ∼10-35% lower than that observed in humans (3.1-3.9; Kong et al. 2012; Sun et al. 2012; Wong et al. 2016; Jónsson et al. 2017). Although no estimates are currently available with regards to the number of spermatogonial stem cell (SSC) divisions in aye-ayes (or any other strepsirrhine), the species will likely incur more frequent SSC divisions than humans (∼23 SSC divisions per year post-puberty; Heller and Clermont 1963; Nielsen et al. 1986; Helgason et al. 2003; and see review by Drost and Lee 1995) due to the shorter spermatogenesis cycle length commonly associated with faster reproduction. Humans reach sexual maturity at the age of ∼13 years (Heller and Clermont 1963); thus, assuming a generation time of 25 years (Fenner 2005), males will have incurred approximately 276 SSC divisions post-puberty at the time of reproduction. In contrast, aye-ayes reach sexual maturity much earlier, at the age of ∼1 year on average (Winn 1994). Assuming a gestation period of 165 days (based on data available in The Animal Ageing and Longevity Database, AnAge) and a higher rate of 33 SSC divisions per year post-puberty (based on available estimates in cercopithecoids; Chowdhury and Steinberger 1976), the number of SSC divisions post-puberty at the time of reproduction in the two sires of the individuals for which the parent-of-origin for the inherited DNMs could be determined is ∼232-260, or about 6–16% fewer than in humans. For comparison, a sexual maturity towards the end of the estimated range (i.e., 36 months; Winn 1994) would yield estimates of ∼166-195 SSC divisions post-puberty, or about 29-40% fewer than in humans.

These considerations may potentially explain both the lower male mutation bias, as well as the similar paternal age effects in young male aye-ayes (with ∼1.2 additional paternal mutations per year of the sire’s age at birth; note that no maternal age effect was observed, likely due to the small sample size; Figure 3c), relative to humans (∼0.9-2.0 additional paternal mutations per year; Conrad et al. 2011; Kong et al. 2012; Besenbacher et al. 2016; Jónsson et al. 2017; and see Table 1 in Moorjani et al. 2016b). Interestingly however, the availability of data across a long reproductive period in aye-ayes (encompassing 15.9 years in males and 19.1 years in females – one of the largest reproductive spans captured in a pedigree-based mutation rate study in primates outside of humans and rhesus macaques to date) demonstrated that the mutation rate in aye-ayes increases much more rapidly with parental age than in humans. For example, consistent with an earlier onset of puberty, aye-ayes exhibit a higher per-site germline mutation rate (∼2.0 x 10^-8^) than humans (∼1.1-1.3 x 10^-8^) at the age of 25 years (Figure 3a). This observation is similar to that of other small-statured primates such as rhesus macaques (Figure 3a), and is likely driven by a combination of a larger number of cell divisions (as expected from the longer reproductive longevity) as well as potentially by other life history traits including the strong male / sperm competition pervasive in polygynandrous mating systems. Despite the overall strong trend, some caution is warranted in such species comparisons, however, as differences across studies in sequencing design (most notably coverage) and computational pipelines can render estimates incomparable (see discussions in Pfeifer 2021; Bergeron et al. 2022).

## CONCLUSION

Our findings underscore the notion that there is no single mutation rate for any given species (see the discussion in Moorjani et al. 2016b), and that data from multiple trios spanning reproductive life is crucial for quantifying variation in the rates and patterns of mutation. For example, in this first detailed look at these mutational dynamics in aye-ayes, we observed amongst the lowest mutation rates in a primate when considering young parents in our pedigree (with maternal and paternal ages at birth of 9.2 and 11.2 years, respectively). Notably though, aye-ayes are thought to reproduce much earlier in the wild, at an average age of 3.5 years (Ross 2003), suggesting rates in natural populations that are potentially even lower, thus likely contributing to the limited genetic diversity characterizing this highly endangered species. However, we also noted a strong parental age effect, with mutation rates increasing much more rapidly with parental age in aye-ayes than in humans, as expected from the greater number of spermatogonial stem cell divisions post-puberty in males. Furthermore, in examining this branch representing a basal split on the primate tree, we observed a mutation spectrum in aye-ayes that is highly similar to those of the much more heavily-studied haplorrhines, likely suggesting a deep evolutionary conservation of the molecular machinery that dictates, at least in part, the rates and patterns of mutation. Given the ever-decreasing cost of sequencing, we anticipate that future studies will continue to illuminate mutational patterns both within- and between-species, and that this more sophisticated characterization of the source of genetic variation will be integrated into existing statistical frameworks in order to gain a better understanding of the evolutionary genomics and chronology of the primate clade (Johri et al. 2022).

## MATERIALS AND METHODS

### Animal subjects

This study was approved by the Duke Lemur Center’s Research Committee (protocol BS-3-22-6) and Duke University’s Institutional Animal Care and Use Committee (protocol A216-20-11), and performed in compliance with all regulations regarding the care and use of captive primates, including the U.S. National Research Council’s Guide for the Care and Use of Laboratory Animals and the U.S. Public Health Service’s Policy on Human Care and Use of Laboratory Animals.

### Whole-genome sequencing

Peripheral blood samples were collected from 14 captive aye-aye (*D. madagascariensis*) individuals from a single three-generation pedigree housed at the Duke Lemur Center (DLC). For each sample, genomic DNA was extracted using the PureLink Genomic DNA Mini Kit (Invitrogen, Waltham, MA, USA), a 150bp paired-end library was prepared using the NEBNext Ultra II DNA PCR-free Library Prep Kit (New England Biolabs, Ipswich, MA, USA), and whole-genome sequenced on the Illumina NovaSeq platform (Illumina, San Diego, CA, USA) to an average coverage of >50X (range: 48.5X to 54.5X per individual; Supplementary Table 1). Figure 1 displays the structure of the pedigree, including the date of birth of the P_0_ and F_1_ individuals in each of the seven trios (i.e., three trios in family 1 and four trios in family 2).

### Data pre-processing

To avoid spurious variant calls, the sequencing data was pre-processed following the guidelines for producing high-quality single nucleotide polymorphism data recommended by Pfeifer (2017b). In brief, raw read data was formatted by marking sequencing adapters using the GATK *MarkIlluminaAdapters* v.4.2.6.1 tool (van der Auwera and O’Connor 2020) and removing bases with quality scores < 20 from the 3’ read-ends using TrimGalore v.0.6.10 (https://github.com/FelixKrueger/TrimGalore). Quality-controlled reads were then mapped to the recently released high-quality, chromosome-level genome assembly for the species (DMad_hybrid; GenBank accession number: JBFSEQ000000000; Versoza and Pfeifer 2024) using BWA-MEM v.0.7.17 (Li and Durbin 2009) with the ’ *-M* ’ option enabled to flag non-primary alignments, and marking duplicates using GATK’s *MarkDuplicates* v.4.2.6.1.

### Variant discovery

Variant discovery followed the GATK Best Practices for non-model organisms (van der Auwera et al. 2013; van der Auwera and O’Connor 2020). Specifically, in the absence of a set of experimentally validated polymorphisms for the species that may be used to identify and correct systematic biases in the sequencing data, an initial round of variant calling was performed from high-quality mappings (’ *--minimum-mapping-quality* 40 ’) of the original (unrecalibrated) data, individual samples merged, and jointly genotyped using GATK’s *HaplotypeCaller* (in ’ *-ERC* GVCF ’ mode with the ’ *--pcr-indel-model* ’ set to NONE as a PCR-free sequencing protocol was used), *CombineGVCFs*, and *GenotypeGVCFs* v.4.2.6.1, respectively. Initial calls were then bootstrapped to create a high-confidence variant dataset for Base Quality Score Recalibration (BQSR) by controlling the transition-transversion ratio, following the methodology described in Auton, Fledel-Alon, Pfeifer, Venn et al. 2012. In brief, GATK’s *SelectVariants* v.4.2.6.1 was used to limit the variant set to biallelic (’ *--restrict-alleles-to* BIALLELIC ’) single nucleotide polymorphisms (’ *--select-type-to-include* SNP’) genotyped in all individuals (’ *AN==*28 ’). Next, variants were removed using BCFtools *filter* v.1.14 (Danecek et al. 2021) using the following "hard filter" criteria with acronyms as defined by GATK: the read depth (*DP*) was less than half or more than twice the genome-wide average, the variant confidence / quality by depth (*QD*) was smaller than 10, the genotype quality (*GQ*) was smaller than 50, the Phred-scaled *p*-value using Fisher’s exact test to detect strand bias (*FS*) was larger than 10, the symmetric odds ratio to detect strand bias (*SOR*) was larger than 1.5, the *Z*-scores from the Wilcoxon rank sum tests of alternative vs. reference read mapping qualities (*MQRankSum*) and position bias (*ReadPosRankSum*) were smaller than -12.5 and -8.0, respectively.

With this high-confidence bootstrapped variant dataset on hand, GATK’s *IndelRealigner* (*RealignerTargetCreator* and *IndelRealigner* v.3.8) and BQSR (*BaseRecalibrator* and *ApplyBQSR* v.4.2.6.1) protocols were applied to the initial read mappings to improve alignments around small insertions and deletions and to correct for systematic errors in base quality (Supplementary Figure 2). A further round of duplicate marking was then performed using GATK’s *MarkDuplicates* v.4.2.6.1, prior to the final round of variant calling and genotyping using the high-quality, realigned, recalibrated data as detailed above but emitting confidence scores at all sites (by using the ’ *-ERC* BP_RESOLUTION ’ mode in the *HaplotypeCaller* and the ’ *-all-sites* ’ flag in the *GenotypeGVCFs* tool) and adjusting the heterozygosity parameter (’ *--heterozygosity* ’) to the species-specific level (*i.e.*, 0.0005; Perry et al. 2013). Lastly, the resulting call set was separated into autosomal biallelic SNPs and monomorphic (*i.e.*, invariant) sites genotyped in all individuals (Supplementary Table 2).

### Identification of DNMs

In order to identify DNMs, the variant dataset was first limited to the 7,907 Mendelian violations observed across the seven trios using BCFtools *view* v.1.14 (Danecek et al. 2021) – that is, sites at which individuals of the P_0_ generation were homozygous for the reference allele ("0/0") and at least one of their focal F_1_ offspring was heterozygous ("0/1" or "1/0"). To increase specificity, Mendelian violations were then re-genotyped using Graphtyper *genotype* v.2.7.2 (Eggertsson et al. 2017), resulting in 459 DNM candidates with high-confidence in the mutant allele that passed built-in sample- and record-level filter. Following the methodology described in Pfeifer (2017a), sequencing reads carrying the DNM candidates were visually inspected for common signs of sequencing, read mapping, variant calling, and/or genotyping errors (for an example, see Figure 4 in Pfeifer 2017b) to eliminate false positives using the Integrated Genomics Viewer (IGV) v.2.16.1 (Thorvaldsdóttir et al. 2012). A total of 323 out of 459 candidate sites passed this manual curation performed independently by two researchers (Supplementary Table 3; IGV screenshots are provided in the Supplementary Materials).

### Sanity checks

As primates generally exhibit low per-site mutation rates (at the order of 10^-9^-10^-8^), few genuine DNMs are expected to be observed in unrelated individuals. To test this, the validated DNMs were screened against segregating variation previously reported in 12 wild aye-aye individuals (Perry et al. 2013). Additionally, as incorrect read mappings can result in spurious variant calls (Pfeifer 2017b), the validated DNMs were also checked for an overlap with regions harboring structural variation (Versoza et al. 2024) which are particularly prone to alignment errors from short-read data due to alterations of the local genomic architecture (Sedlazeck et al. 2018). Lastly, based on Mendel’s principles of segregation (Mendel 1866), the expectation for transmission of a genuine DNM to the next generation is 50%. A Fisher’s exact test was performed in R v.4.2.2 (R Core Team 2022) to assess whether the transmission rates from the F_1_ individuals to their F_2_ offspring were consistent with this expectation.

### Annotation of DNMs

Validated DNMs were annotated using ANNOVAR v.2020-06-08 (Wang et al. 2020) to categorize them by genomic region (i.e., intergenic, upstream, exonic, exonic non-coding RNA [ncRNA], intronic, intronic ncRNA, 3’ and 5’ UTR, and downstream) based on the annotations available for the aye-aye genome assembly (DMad_hybrid; GenBank accession number: JBFSEQ010000000; Versoza and Pfeifer 2024). Additionally, to obtain a baseline expectation for the distribution, the pipeline was also run on the complete call set of autosomal biallelic SNPs and monomorphic sites genotyped in all individuals (Supplementary Table 2). The genomic distribution was plotted using ggplot2 v.3.4.1 (Wickham 2016) in R v.4.2.2 (R Core Team 2022) and a χ^2^-test was performed to compare the proportion of DNMs in each genomic region against the overall genome-wide composition.

### Characterization of primate mutation spectra

DNMs were grouped by mutation type – that is A>C, A>G, A>T, C>A, and C>G mutations as well as C>T transitions that occurred within a CpG context (i.e., CpG>TpG) and outside of a CpG context (i.e., CpH>TpH), with reverse complements collapsed – based on the aye-aye genome assembly (DMad_hybrid; GenBank accession number: JBFSEQ010000000; Versoza and Pfeifer 2024), with the relative proportion of each mutation type representing the mutation spectrum. The mutation spectrum of aye-ayes was compared with those of haplorrhines – as assessed from catarrhines (humans [Kong et al. 2012; Besenbacher et al. 2016; Goldmann et al. 2016; Rahbari et al. 2016], chimpanzees [Venn et al. 2014; Besenbacher et al. 2019], gorillas [Besenbacher et al. 2019], orangutans [Besenbacher et al. 2019], vervet monkeys [Pfeifer 2017a], rhesus macaques [Wang et al. 2020], and baboons [Wu et al. 2020]) and platyrrhines (owl monkeys [Thomas et al. 2018]). Mutational spectra were plotted in R v.4.2.2 (R Core Team 2022) using code provided by Gregg Thomas (https://github.com/gwct/owl-monkey), and a χ^2^-test was performed to compare the mutation spectra for the eight primate species.

### Clustering of DNMs

In order to identify non-random clustering of DNMs, DNMs were analyzed using VCFtools v.0.1.14 (Danecek et al. 2011) in windows (’ --SNPdensity ’) of size 1kb, 10kb, 100kb, and 1Mb.

### Estimation of the false negative rate

Following Pfeifer (2017a), the false negative rate of the experiment was estimated based on simulations of synthetic DNMs that were "spiked" into the sequencing reads. As accurate haplotype resolution is important for the discovery of genetic variants, the DNMs were simulated in the focal offspring in a haplotype-aware manner. In brief, WhatsHap *phase* v.2.3 (Patterson et al. 2015; Garg et al. 2016) was used in the pedigree-aware mode (’ *--ped* ’) to phase reads by combining read-based phasing with phasing based on the Mendelian rules of inheritance, assuming a constant recombination rate of ∼1 cM/Mb across the genome as previously observed in the species (Versoza, Lloret-Villas et al. 2024). Next, the *addsnv.py* script included in BAMSurgeon v.1.4.1 (Ewing et al. 2015) was used to add 1,000 DNMs at random in the haplotype-resolved reads of the F_1_ individuals, setting the maximum allowable minor allele frequency of linked SNPs to 0.1 (’ -*s* 0.1 ’). With this setting, BAMSurgeon successfully added 684 synthetic DNMs that mimicked the allele balance observed at genuine heterozygous sites in the trios (i.e., sites were one of the parents was homozygous for the reference allele, the other parent homozygous for the alternative allele, and their joint offspring heterozygous; Supplementary Figure 3). Reads were analyzed following the protocols described in "Variant discovery" and "Identification of DNMs". In total, GATK identified 1,449 Mendelian violations – a 2-fold excess of the number of synthetic DNMs added (note that GATK correctly identified all synthetic DNMs as non-reference alleles though one DNM was classified as an insertion rather than a SNP). Re-genotyping these Mendelian violations with Graphtyper discovered 619 out of the 684 synthetic DNMs (no additional Mendelian violations were present in the re-genotyped dataset), yielding a false negative rate of 9.5%.

### Phasing of DNMs

In order to determine the parent-of-origin of the DNMs, a combination of direct (read-based) and indirect (genetic) phasing was applied. First, WhatsHap v.2.3 (Patterson et al. 2015; Garg et al. 2016) was used to phase reads from all individuals as described in "Estimation of the false negative rate". Additionally, DNMs carried by the two F_1_ individuals with multiple offspring (i.e., individuals 7 and 8) were phased by transmission to their offspring in the F_2_ generation (i.e., individuals 12, 13, and 14). In brief, the three-generation pedigree data allowed for the phasing of variants through "phase-informative" markers – that is, sites at which the P_0_ individuals have distinct genotypes, the focal F_1_ individual is heterozygous, and either the F_1_’s partner or their joint F_2_ offspring is homozygous (for a schematic representation, see Figure 1b in Versoza, Weiss et al. 2024). Using such phase-informative markers, the parent-of-origin of the DNM transmitted to the third (F_2_) generation can then be established from the phase of the haplotype block. Haplotype blocks were required to be at least 0.5 Mb in length and contain a minimum of 100 phase-informative markers. DNMs with incongruous haplotype phase between F_2_ siblings were classified as ambiguous and thus not assigned a parental haplotype. Through this approach, 10 and 27 DNMs were assigned as maternal and paternal in origin, respectively.

## ACKNOWLEDGEMENTS

We would like to thank the Duke Lemur Center for providing the aye-aye samples used in this study. DNA extraction, library preparation, and Illumina sequencing were conducted at Azenta Life Sciences (South Plainfield, NJ, USA). Computations were performed on the Sol supercomputer at Arizona State University (Jennewein et al. 2023). This is Duke Lemur Center publication # XXXX.

## REFERENCES

Auton A, Fledel-Alon A, Pfeifer S, Venn O, Ségurel L, Street T, Leffler EM, Bowden R, Aneas I, Broxholme J, et al. 2012. A fine-scale chimpanzee genetic map from population sequencing. Science. 336(6078): 193–198.

Baer CF, Miyamoto MM, Denver DR. 2007. Mutation rate variation in multicellular eukaryotes: causes and consequences. Nat Rev Genet. 8(8): 619–631.

Beal MA, Glenn TC, Somers CM. 2012. Whole genome sequencing for quantifying germline mutation frequency in humans and model species: cautious optimism. Mutat Res. 750(2): 96–106.

Beichman AC, Zhu L, Harris K. 2024. The evolutionary interplay of somatic and germline mutation rates. Annu Rev Biomed Data Sci. 7(1): 83–105.

Bergeron LA, Besenbacher S, Bakker J, Zheng J, Li P, Pacheco G, Sinding MS, Kamilari M, Gilbert MTP, Schierup MH, et al. 2021. The germline mutational process in rhesus macaque and its implications for phylogenetic dating. Giga Science. 10(5): giab029.

Bergeron LA, Besenbacher S, Turner T, Versoza CJ, Wang RJ, Price AL, Armstrong E, Riera M, Carlson J, Chen HY, et al. 2022. The Mutationathon highlights the importance of reaching standardization in estimates of pedigree-based germline mutation rates. Elife. 11: e73577.

Bergeron LA, Besenbacher S, Zheng J, Li P, Bertelsen MF, Quintard B, Hoffman JI, Li Z, St Leger J, Shao C, et al. 2023. Evolution of the germline mutation rate across vertebrates. Nature. 615(7951): 285–291.

Besenbacher S, Hvilsom C, Marques-Bonet T, Mailund T, Schierup MH. 2019. Direct estimation of mutations in great apes reconciles phylogenetic dating. Nat Ecol Evol. 3(2): 286–292.

Besenbacher S, Sulem P, Helgason A, Helgason H, Kristjansson H, Jonasdottir A, Jonasdottir A, Magnusson OT, Thorsteinsdottir U, Masson G, et al. 2016. Multi-nucleotide *de novo* mutations in humans. PLoS Genet. 12(11): e1006315.

Biesecker LG, Spinner NB. 2013. A genomic view of mosaicism and human disease. Nat Rev Genet. 14(5): 307–320.

Campbell CD, Chong JX, Malig M, Ko A, Dumont BL, Han L, Vives L, O’Roak BJ, Sudmant PH, Shendure J, et al. 2012. Estimating the human mutation rate using autozygosity in a founder population. Nat Genet. 44(11): 1277–1281.

Campbell CR, Tiley GP, Poelstra JW, Hunnicutt KE, Larsen PA, Lee HJ, Thorne JL, Dos Reis M, Yoder AD. 2021. Pedigree-based and phylogenetic methods support surprising patterns of mutation rate and spectrum in the gray mouse lemur. Heredity (Edinb). 127(2): 233–244.

Chimpanzee Sequencing and Analysis Consortium. 2005. Initial sequence of the chimpanzee genome and comparison with the human genome. Nature. 437(7055): 69–87.

Chintalapati M, Moorjani P. 2020. Evolution of the mutation rate across primates. Curr Opin Genet Dev. 62: 58–64.

Chowdhury AK, Steinberger E. 1976. A study of germ cell morphology and duration of spermatogenic cycle in the baboon, *Papio anubis*. Anat Rec. 185(2): 155–169.

Conrad DF, Keebler JE, DePristo MA, Lindsay SJ, Zhang Y, Casals F, Idaghdour Y, Hartl CL, Torroja C, Garimella KV, et al. 2011. Variation in genome-wide mutation rates within and between human families. Nat Genet. 43(7): 712–714.

Crow JF. 2000. The origins, patterns and implications of human spontaneous mutation. Nat Rev Genet. 1(1): 40–47.

Crow JF. 2006. Age and sex effects on human mutation rates: an old problem with new complexities. J Radiat Res. 47(Suppl B): B75–B82.

Danecek P, Auton A, Abecasis G, Albers CA, Banks E, DePristo MA, Handsaker RE, Lunter G, Marth GT, Sherry ST, et al. 2011. The variant call format and VCFtools. Bioinformatics. 27(15): 2156–2158.

Danecek P, Bonfield JK, Liddle J, Marshall J, Ohan V, Pollard MO, Whitwham A, Keane T, McCarthy SA, Davies RM, 2021. Twelve years of SAMtools and BCFtools. Giga Science. 10(2): giab008.

Drake JW, Charlesworth B, Charlesworth D, Crow JF. 1998. Rates of spontaneous mutation. Genetics. 148(4): 1667–1686.

Driscoll DJ, Migeon BR. 1990. Sex difference in methylation of single-copy genes in human meiotic germ cells: implications for X chromosome inactivation, parental imprinting, and origin of CpG mutations. Somat Cell Mol Genet. 16(3): 267–282.

Drost JB, Lee WR. 1995. Biological basis of germline mutation: comparisons of spontaneous germline mutation rates among drosophila, mouse, and human. Environ Mol Mutagen. 25(Suppl 26): 48–64.

Eggertsson HP, Jonsson H, Kristmundsdottir S, Hjartarson E, Kehr B, Masson G, Zink F, Hjorleifsson KE, Jonasdottir A, Jonasdottir A, et al. 2017. Graphtyper enables population-scale genotyping using pangenome graphs. Nat Genet. 49(11): 1654–1660.

Ellegren H. 2007. Characteristics, causes and evolutionary consequences of male-biased mutation. Proc Biol Sci. 274(1606): 1–10.

Ewing AD, Houlahan KE, Hu Y, Ellrott K, Caloian C, Yamaguchi TN, Bare JC, P’ng C, Waggott D, Sabelnykova VY, et al. 2015. Combining tumor genome simulation with crowdsourcing to benchmark somatic single-nucleotide-variant detection. Nat Methods. 12(7): 623–630.

Fenner JN. 2005. Cross-cultural estimation of the human generation interval for use in genetics-based population divergence studies. Am J Phys Anthropol. 128(2): 415–423.

Francioli LC, Polak PP, Koren A, Menelaou A, Chun S, Renkens I; Genome of the Netherlands Consortium; van Duijn CM, Swertz M, Wijmenga C, et al. 2015. Genome-wide patterns and properties of *de novo* mutations in humans. Nat Genet. 47(7): 822–826.

Garg S, Martin M, Marschall T. 2016. Read-based phasing of related individuals. Bioinformatics. 32(12): i234–i242.

Gao Z, Moorjani P, Sasani TA, Pedersen BS, Quinlan AR, Jorde LB, Amster G, Przeworski M. 2019. Overlooked roles of DNA damage and maternal age in generating human germline mutations. Proc Natl Acad Sci U S A. 116(19): 9491–9500.

Gao Z, Wyman MJ, Sella G, Przeworski M. 2016. Interpreting the dependence of mutation rates on age and time. PLoS Biol. 14(1): e1002355.

Goldmann JM, Wong WS, Pinelli M, Farrah T, Bodian D, Stittrich AB, Glusman G, Vissers LE, Hoischen A, Roach JC, et al. 2016. Parent-of-origin-specific signatures of *de novo* mutations. Nat Genet. 48(8): 935–939.

Goriely A. 2016. Decoding germline *de novo* point mutations. Nat Genet. 48(8): 823–824.

Haldane JB. 1935. The rate of spontaneous mutation of a human gene. J Genet. 83(3): 235–244.

Haldane JB. 1947. The mutation rate of the gene for haemophilia, and its segregation ratios in males and females. Ann Eugen. 13(4): 262–271.

Helgason A, Hrafnkelsson B, Gulcher JR, Ward R, Stefánsson K. 2003. A populationwide coalescent analysis of Icelandic matrilineal and patrilineal genealogies: evidence for a faster evolutionary rate of mtDNA lineages than Y chromosomes. Am J Hum Genet. 72(6): 1370–1388.

Heller CG, Clermont Y. 1963. Spermatogenesis in man: an estimate of its duration. Science. 140(3563): 184–186.

Hodgkinson A, Eyre-Walker A. 2011. Variation in the mutation rate across mammalian genomes. Nat Rev Genet. 12(11): 756–766.

Hwang DG, Green P. 2004. Bayesian Markov chain Monte Carlo sequence analysis reveals varying neutral substitution patterns in mammalian evolution. Proc Natl Acad Sci U S A. 101(39): 13994–14001.

Jennewein DM, Lee J, Kurtz C, Dizon W, Shaeffer I, Chapman A, Chiquete A, Burks J, Carlson A, Mason N, et al. 2023. The Sol Supercomputer at Arizona State University. In Practice and Experience in Advanced Research Computing 2023: Computing for the Common Good (PEARC ’23). Association for Computing Machinery, New York, NY, USA, 296–301.

Jónsson H, Sulem P, Arnadottir GA, Pálsson G, Eggertsson HP, Kristmundsdottir S, Zink F, Kehr B, Hjorleifsson KE, Jensson BÖ, et al. 2018. Multiple transmissions of *de novo* mutations in families. Nat Genet. 50(12): 1674–1680.

Jónsson H, Sulem P, Kehr B, Kristmundsdottir S, Zink F, Hjartarson E, Hardarson MT, Hjorleifsson KE, Eggertsson HP, Gudjonsson SA, et al. 2017. Parental influence on human germline *de novo* mutations in 1,548 trios from Iceland. Nature. 549(7673): 519– 522.

Johri P, Aquadro CF, Beaumont M, Charlesworth B, Excoffier L, Eyre-Walker A, Keightley PD, Lynch M, McVean G, Payseur BA, et al. 2022. Recommendations for improving statistical inference in population genomics. PLoS Biol. 20(5): e3001669.

Kessler MD, Loesch DP, Perry JA, Heard-Costa NL, Taliun D, Cade BE, Wang H, Daya M, Ziniti J, Datta S, et al. 2020. *De novo* mutations across 1,465 diverse genomes reveal mutational insights and reductions in the Amish founder population. Proc Natl Acad Sci U S A. 117(5): 2560–2569.

Kimura M. 1968. Evolutionary rate at the molecular level. Nature. 217(5129): 624–626.

Kimura M. 1983. The neutral theory of molecular evolution. Cambridge Univ Press, Cambridge, MA.

Kondrashov AS. 2003. Direct estimates of human per nucleotide mutation rates at 20 loci causing Mendelian diseases. Hum Mutat. 21(1): 12–27.

Kong A, Frigge ML, Masson G, Besenbacher S, Sulem P, Magnusson G, Gudjonsson SA, Sigurdsson A, Jonasdottir A, Jonasdottir A, et al. 2012. Rate of *de novo* mutations and the importance of father’s age to disease risk. Nature. 488(7412): 471–475.

Kuderna LFK, Gao H, Janiak MC, Kuhlwilm M, Orkin JD, Bataillon T, Manu S, Valenzuela A, Bergman J, Rousselle M, et al. 2023. A global catalog of whole-genome diversity from 233 primate species. 380(6648): 906–913.

Leffler EM, Gao Z, Pfeifer S, Ségurel L, Auton A, Venn O, Bowden R, Bontrop R, Wall JD, Sella G, et al. 2013. Multiple instances of ancient balancing selection shared between humans and chimpanzees. Science. 339(6127): 1578–1582.

Li H, Durbin R. 2009. Fast and accurate short read alignment with Burrows-Wheeler transform. Bioinformatics. 25(14): 1754–1760.

Louis EE, Sefczek TM, Randimbiharinirina DR, Raharivololona B, Rakotondrazandry JN, Manjary D, Aylward M, Ravelomandrato F. 2020. Daubentonia madagascariensis. The IUCN Red List of Threatened Species 2020: e.T6302A115560793.

Lynch M. 2010. Rate, molecular spectrum, and consequences of human mutation. Proc Natl Acad Sci U S A. 107(3): 961–968.

Lynch M, Ackerman MS, Gout JF, Long H, Sung W, Thomas WK, Foster PL. 2016. Genetic drift, selection and the evolution of the mutation rate. Nat Rev Genet. 17(11): 704–714.

Mendel G. 1866. Versuche über Pflanzen-Hybriden. Verhandlungen des Naturforschenden Vereines, Abhandlungern, Brünn. 4: 3–47.

Michaelson JJ, Shi Y, Gujral M, Zheng H, Malhotra D, Jin X, Jian M, Liu G, Greer D, Bhandari A, et al. 2012. Whole-genome sequencing in autism identifies hot spots for *de novo* germline mutation. Cell. 151(7): 1431–1442.

Mittermeier RA, Hawkins F, Louis EE. 2010. Lemurs of Madagascar. 3rd ed. Conservation International.

Mohrenweiser HW, Wilson DM 3rd, Jones IM. 2003. Challenges and complexities in estimating both the functional impact and the disease risk associated with the extensive genetic variation in human DNA repair genes. Mutat Res. 526(1-2): 93–125.

Moorjani P, Amorim CE, Arndt PF, Przeworski M. 2016a. Variation in the molecular clock of primates. Proc Natl Acad Sci U S A. 113(38): 10607–10612.

Moorjani P, Gao Z, Przeworski M. 2016b. Human germline mutation and the erratic evolutionary clock. PLoS Biol. 14(10): e2000744.

Nachman MW. 2004. Haldane and the first estimates of the human mutation rate. J Genet. 83(3): 231–233.

Nachman MW, Crowell SL. 2000. Estimate of the mutation rate per nucleotide in humans. Genetics. 156(1): 297–304.

Nielsen CT, Skakkebaek NE, Richardson DW, Darling JA, Hunter WM, Jørgensen M, Nielsen A, Ingerslev O, Keiding N, Müller J. 1986. Onset of the release of spermatozoa (spermarche) in boys in relation to age, testicular growth, pubic hair, and height. J Clin Endocrinol Metab. 62(3): 532–535.

Nielsen R, Akey JM, Jakobsson M, Pritchard JK, Tishkoff S, Willerslev E. 2017. Tracing the peopling of the world through genomics. Nature. 541(7637): 302–310.

Patterson M, Marschall T, Pisanti N, van Iersel L, Stougie L, Klau GW, Schönhuth A. 2015. WhatsHap: weighted haplotype assembly for future-generation sequencing reads. J Comput Biol. 22(6): 498–509.

Perry GH, Louis EE Jr, Ratan A, Bedoya-Reina OC, Burhans RC, Lei R, Johnson SE, Schuster SC, Miller W. 2013. Aye-aye population genomic analyses highlight an important center of endemism in northern Madagascar. Proc Natl Acad Sci U S A. 110(15): 5823–5828.

Pfeifer SP. 2017a. Direct estimate of the spontaneous germ line mutation rate in African green monkeys. Evolution. 71(12): 2858–2870.

Pfeifer SP. 2017b. From next-generation resequencing reads to a high-quality variant data set. Heredity (Edinb). 118(2): 111–124.

Pfeifer SP. 2020. Spontaneous mutation rates. In Ho SYW (ed) The Molecular Evolutionary Clock. Theory and Practice. Springer Nature, pp. 35–44.

Pfeifer SP. 2021. Studying mutation rate evolution in primates-the effects of computational pipelines and parameter choices. Giga Science. 10(10):giab069.

R Core Team. 2022. R: a language and environment for statistical computing. R Foundation for Statistical Computing, Vienna, Austria. https://www.R-project.org/.

Rahbari R, Wuster A, Lindsay SJ, Hardwick RJ, Alexandrov LB, Turki SA, Dominiczak A, Morris A, Porteous D, Smith B, et al. 2016. Timing, rates and spectra of human germline mutation. Nat Genet. 48(2): 126–133.

Randimbiharinirina DR, Sefczek TM, Raharivololona BM, Rostant Andriamalala Y, Ratsimbazafy J, Louis Jr EE. 2019. Aye-aye *Daubentonia madagascariensis* (Gmelin, 1788). In: Primates in Peril: The World’s 25 Most Endangered Primates 2018–2020.

Schwitzer, C et al., editors. Washington, DC, USA.: IUCN SSC Primate Specialist Group (PSG), International Primatological Society (IPS), Global Wildlife Conservation (GWC), Bristol Zoological Society (BZS).

Roach JC, Glusman G, Smit AF, Huff CD, Hubley R, Shannon PT, Rowen L, Pant KP, Goodman N, Bamshad M, et al. 2010. Analysis of genetic inheritance in a family quartet by whole-genome sequencing. Science. 328(5978): 636–639.

Ross C. 2003. Life history, infant care strategies, and brain size in primates. In: Kappeler PM, Pereira ME, editors. Primate life histories and socioecology. Chicago (IL): Chicago University Press. pp. 266–284.

Samuels ME, Friedman JM. 2015. Genetic mosaics and the germ line lineage. Genes (Basel). 6(2): 216–237.

Sasani TA, Pedersen BS, Gao Z, Baird L, Przeworski M, Jorde LB, Quinlan AR. 2019. Large, three-generation human families reveal post-zygotic mosaicism and variability in germline mutation accumulation. Elife. 8:e46922.

Scally A. 2016. Mutation rates and the evolution of germline structure. Philos Trans R Soc Lond B Biol Sci. 371(1699): 20150137.

Sedlazeck FJ, Rescheneder P, Smolka M, Fang H, Nattestad M, von Haeseler A, Schatz MC. 2018. Accurate detection of complex structural variations using single-molecule sequencing. Nat Methods. 15(6): 461–468.

Ségurel L, Wyman MJ, Przeworski M. 2014. Determinants of mutation rate variation in the human germline. Annu Rev Genomics Hum Genet. 15: 47–70.

Shendure J, Akey JM. 2015. The origins, determinants, and consequences of human mutations. Science. 349(6255): 1478–1483.

Slotkin RK, Martienssen R. 2007. Transposable elements and the epigenetic regulation of the genome. Nat Rev Genet. 8(4): 272–285.

Soni V, Terbot JW, Versoza CJ, Pfeifer SP, Jensen JD. 2024. A whole-genome scan for evidence of recent positive and balancing selection in aye-ayes utilizing a well-fit evolutionary baseline model. In preprint. BioRxiv.

Sun JX, Helgason A, Masson G, Ebenesersdóttir SS, Li H, Mallick S, Gnerre S, Patterson N, Kong A, Reich D, et al. 2012. A direct characterization of human mutation based on microsatellites. Nat Genet. 44(10): 1161–1165.

Sung W, Ackerman MS, Miller SF, Doak TG, Lynch M. 2012. Drift-barrier hypothesis and mutation-rate evolution. Proc Natl Acad Sci U S A. 109(45): 18488–18492.

Suzzi-Simmons A. 2023. Status of deforestation of Madagascar. Glob Ecol Conserv. 42:e02389.

Tatsumoto S, Go Y, Fukuta K, Noguchi H, Hayakawa T, Tomonaga M, Hirai H, Matsuzawa T, Agata K, Fujiyama A. 2017. Direct estimation of *de novo* mutation rates in a chimpanzee parent-offspring trio by ultra-deep whole genome sequencing. Sci Rep. 7(1): 13561.

Terbot JW, Soni V, Versoza CJ, Pfeifer SP, Jensen JD. 2024. Inferring the demographic history of aye-ayes (*Daubentonia madagascariensis)* from high-quality, whole-genome, population-level data. In preprint. BioRxiv.

Thomas GWC, Wang RJ, Puri A, Harris RA, Raveendran M, Hughes DST, Murali SC, Williams LE, Doddapaneni H, Muzny DM, et al. 2018. Reproductive longevity predicts mutation rates in primates. Curr Biol. 28(19): 3193–3197.

Thorvaldsdóttir H, Robinson JT, Mesirov JP. 2013. Integrative Genomics Viewer (IGV): high-performance genomics data visualization and exploration. Brief Bioinform. 14(2): 178–192.

Tran LAP, Pfeifer SP. 2018. Germline mutation rates in Old World monkeys. eLS, John Wiley & Sons, Ltd: Chichester.

Tyekucheva S, Makova KD, Karro JE, Hardison RC, Miller W, Chiaromonte F. 2008. Human-macaque comparisons illuminate variation in neutral substitution rates. Genome Biol. 9(4): R76.

van der Auwera GA, Carneiro MO, Hartl C, Poplin R, Del Angel G, Levy-Moonshine A, Jordan T, Shakir K, Roazen D, Thibault J, et al. 2013. From FastQ data to high confidence variant calls: the Genome Analysis Toolkit best practices pipeline. Curr Protoc Bioinformatics. 43(1110): 11.10.1–11.10.33.

van der Auwera GA, O’Connor BD. 2020. Genomics in the cloud: using Docker, GATK, and WDL in Terra. Sebastopol: O’Reilly Media.

Venn O, Turner I, Mathieson I, de Groot N, Bontrop R, McVean G. 2014. Strong male bias drives germline mutation in chimpanzees. Science. 344(6189): 1272–1275.

Versoza CJ, Jensen JD, Pfeifer SP. 2024. The landscape of structural variation in aye-ayes (*Daubentonia madagascariensis)*. In preprint. BioRxiv.

Versoza CJ, Lloret-Villas A, Jensen JD, Pfeifer SP. 2024. A pedigree-based map of crossovers and non-crossovers in aye-ayes (*Daubentonia madagascariensis)*. In preprint. BioRxiv.

Versoza CJ, Pfeifer SP. 2024. A hybrid genome assembly of the endangered aye-aye (*Daubentonia madagascariensis*). G3 (Bethesda). 14(10): jkae185.

Versoza CJ, Weiss S, Johal R, La Rosa B, Jensen JD, Pfeifer SP. 2024. Novel insights into the landscape of crossover and noncrossover events in rhesus macaques (*Macaca mulatta*). Genome Biol Evol. 16(1): evad223.

Wang H, Zhu X. 2014. *De novo* mutations discovered in 8 Mexican American families through whole genome sequencing. BMC Proc. 8(Suppl 1):S24.

Wang K, Li M, Hakonarson H. 2010. ANNOVAR: functional annotation of genetic variants from high-throughput sequencing data. Nucleic Acids Res. 38(16): e164.

Wang RJ, Thomas GWC, Raveendran M, Harris RA, Doddapaneni H, Muzny DM, Capitanio JP, Radivojac P, Rogers J, Hahn MW. 2020. Paternal age in rhesus macaques is positively associated with germline mutation accumulation but not with measures of offspring sociability. Genome Res. 30(6): 826–834.

Wickham H. 2016. ggplot2: elegant graphics for data analysis. Springer-VerlagNew York.

Wilson Sayres MA, Venditti C, Pagel M, Makova KD. 2011. Do variations in substitution rates and male mutation bias correlate with life-history traits? A study of 32 mammalian genomes. Evolution. 65(10): 2800–2815.

Winn RM. 1994. Development of behaviour in a young aye-aye (*Daubentonia madagascariensis*) in captivity. Folia Primatol (Basel). 62(1-3): 93–107.

Wong WS, Solomon BD, Bodian DL, Kothiyal P, Eley G, Huddleston KC, Baker R, Thach DC, Iyer RK, Vockley JG, et al. 2016. New observations on maternal age effect on germline *de novo* mutations. Nat Commun. 7: 10486.

Woodruff RC, Thompson JNJ. 1992. Have premeiotic clusters of mutation been overlooked in evolutionary theory? J. Evol. Biol. 5:457–646

Wu FL, Strand AI, Cox LA, Ober C, Wall JD, Moorjani P, Przeworski M. 2020. A comparison of humans and baboons suggests germline mutation rates do not track cell divisions. PLoS Biol. 18(8): e3000838.

Yang C, Zhou Y, Marcus S, Formenti G, Bergeron LA, Song Z, Bi X, Bergman J, Rousselle MMC, Zhou C, Zhou L, et al. 2021. Evolutionary and biomedical insights from a marmoset diploid genome assembly. Nature. 594(7862): 227–233.

Zehr SM, Roach RG, Haring D, Taylor J, Cameron FH, Yoder AD. 2014. Life history profiles for 27 strepsirrhine primate taxa generated using captive data from the Duke Lemur Center. Sci Data. 1: 140019.

