## Supplementary Materials for "Characterizing the rates and patterns of *de novo* germline mutations in the aye-aye (*Daubentonia madagascariensis*)"

**Supplementary Table 1**. Sample information, including the sex and date of birth of the parental ($P_0$) and focal ($F_1$) individuals (highlighted in green) as well as their sequencing coverages.

| pedigree ID | sex | sire ID | dam ID | date of birth (yyyy-mm-dd) | sire's age at birth (in days) | dam's age at birth (in days) | coverage |
|---|---|---|---|---|---|---|---|
| 1 | M | – | – | 1986-12-19 | – | – | 50.5 |
| 2 | F | – | – | 1988-11-30 | – | – | 50.2 |
| 3 | M | – | – | 1985-12-18 | – | – | 53.7 |
| 4 | F | – | – | 1983-12-04 | – | – | 52.5 |
| 5 | M | 1 | 2 | 2005-02-22 | 6,640 | 5,928 | 54.1 |
| 6 | M | 1 | 2 | 1998-02-08 | 4,069 | 3,357 | 53.7 |
| 7 | M | 3 | 4 | 1994-06-05 | 3,091 | 3,836 | 54.1 |
| 8 | F | 1 | 2 | 1996-04-15 | 3,405 | 2,693 | 54.5 |
| 9 | M | 3 | 4 | 2010-05-23 | 8,922 | 9,667 | 51.8 |
| 10 | F | 3 | 4 | 2005-12-30 | 7,317 | 8,062 | 52.7 |
| 11 | F | 3 | 4 | 2001-07-30 | 5,703 | 6,448 | 52.0 |
| 12 | F | 7 | 8 | – | – | – | 48.5 |
| 13 | F | 7 | 8 | – | – | – | 52.4 |
| 14 | F | 9 | 8 | – | – | – | 52.1 |

**Supplementary Table 2**. Summary of the variant call set. A total of 3,594,088 autosomal, biallelic, single nucleotide polymorphisms (SNPs) with a transition-transversion ratio (Ts/Tv) of 2.47 were discovered in the accessible genome.

| scaffold | length | # invariant sites | # SNPs | % accessible | Ts/Tv |
|---|---|---|---|---|---|
| 1 | 316,165,773 | 312,369,144 | 499,552 | 98.96 | 2.36 |
| 2 | 290,592,686 | 286,926,937 | 445,239 | 98.89 | 2.39 |
| 3 | 261,424,170 | 258,279,175 | 430,647 | 98.96 | 2.33 |
| 4 | 219,686,500 | 217,268,884 | 340,010 | 99.05 | 2.39 |
| 5 | 215,448,047 | 212,923,711 | 326,720 | 98.98 | 2.45 |
| 6 | 204,016,426 | 201,827,142 | 309,052 | 99.08 | 2.46 |
| 7 | 199,604,927 | 196,630,522 | 327,919 | 98.67 | 2.39 |
| 8 | 162,769,830 | 160,801,230 | 252,071 | 98.95 | 2.41 |
| 10 | 114,896,738 | 113,523,171 | 170,024 | 98.95 | 2.49 |
| 11 | 102,076,017 | 100,745,800 | 154,928 | 98.85 | 2.61 |
| 12 | 67,301,774 | 65,825,151 | 118,802 | 97.98 | 2.45 |
| 13 | 62,733,483 | 61,766,335 | 106,242 | 98.63 | 2.64 |
| 14 | 34,254,822 | 33,756,217 | 63,096 | 98.73 | 2.56 |
| 15 | 28,257,198 | 27,963,532 | 49,786 | 99.14 | 2.65 |
| ∑ or ∅ | **2,279,228,391** | **2,250,606,951** | **3,594,088** | **98.84** | **2.47** |

**Supplementary Table 3**. *De novo* mutations detected across the seven parent-offspring trios. Locations (*scaffold* and *position*) and *genomic annotation* are defined according to the DMad_hybrid genome assembly. *REF* and *ALT* indicate the reference and alternative alleles, respectively. *Repeat status* indicates whether the mutation is located within a repetitive region of the genome. *CpG>TpG status* indicates C-to-T transitions at CpG sites. *Parental origin* indicates the phase inferred genetic phasing (if available).

| scaffold | position | REF | ALT | recipient ($F_1$) | transmitted ($F_2$) | parental origin | repeat status | CpG>TpG status | genomic annotation |
|---|---|---|---|---|---|---|---|---|---|
| scaffold 1 | 3,782,319 | A | G | 9 | | | 0 | 0 | ncRNA intronic |
| scaffold 1 | 12,054,128 | C | T | 5 | | | 0 | 0 | intronic |
| scaffold 1 | 14,183,338 | C | T | 6 | | | 0 | 0 | intergenic |
| scaffold 1 | 34,169,345 | A | G | 9 | | | 1 | 0 | intergenic |
| scaffold 1 | 51,159,485 | C | T | 10 | | | 1 | 0 | intergenic |
| scaffold 1 | 69,175,149 | T | C | 5 | | | 0 | 0 | intronic |
| scaffold 1 | 84,192,234 | C | T | 10 | | | 0 | 0 | intergenic |
| scaffold 1 | 97,611,037 | C | G | 9 | 14 | | 0 | 0 | intergenic |
| scaffold 1 | 103,338,297 | A | G | 5 | | | 1 | 0 | intergenic |
| scaffold 1 | 109,819,319 | A | G | 8 | 12,14 | paternal | 1 | 0 | intronic |
| scaffold 1 | 111,235,804 | A | G | 11 | | | 0 | 0 | intronic |
| scaffold 1 | 115,417,966 | G | A | 5 | | | 1 | 0 | intergenic |
| scaffold 1 | 119,156,328 | T | C | 9 | | | 0 | 0 | intronic |
| scaffold 1 | 128,603,832 | G | A | 9 | | | 1 | 0 | intergenic |
| scaffold 1 | 130,555,992 | C | T | 6 | | | 0 | 1 | ncRNA intronic |
| scaffold 1 | 137,376,770 | T | C | 10 | | | 0 | 0 | intronic |
| scaffold 1 | 142,152,454 | G | T | 9 | 14 | | 0 | 0 | intergenic |
| scaffold 1 | 143,217,359 | G | A | 5 | | | 0 | 0 | intronic |
| scaffold 1 | 144,285,461 | G | A | 6,8 | 13 | maternal | 0 | 1 | intergenic |
| scaffold 1 | 152,524,873 | T | A | 9 | | | 1 | 0 | intergenic |
| scaffold 1 | 155,744,337 | C | A | 9 | | | 0 | 0 | intergenic |
| scaffold 1 | 161,685,074 | T | C | 7 | 12 | paternal | 1 | 0 | intergenic |
| scaffold 1 | 163,542,285 | G | C | 11 | | | 0 | 0 | intronic |
| scaffold 1 | 170,273,061 | C | A | 6 | | | 0 | 0 | intronic |
| scaffold 1 | 172,641,223 | A | G | 10 | | | 0 | 0 | intergenic |
| scaffold 1 | 176,098,461 | T | C | 10 | | | 0 | 0 | intergenic |
| scaffold 1 | 176,419,328 | C | T | 9 | 14 | | 1 | 0 | intergenic |
| scaffold 1 | 181,469,167 | C | T | 10 | | | 1 | 1 | intergenic |
| scaffold 1 | 186,371,192 | A | T | 11 | | | 0 | 0 | ncRNA intronic |
| scaffold 1 | 190,670,770 | G | A | 10 | | | 0 | 1 | UTR3 |
| scaffold 1 | 200,895,173 | C | T | 11 | | | 0 | 1 | intronic |
| scaffold 1 | 202,516,631 | G | A | 9 | 14 | | 0 | 0 | intronic |
| scaffold 1 | 214,210,022 | A | G | 7 | 12 | maternal | 1 | 0 | intergenic |
| scaffold 1 | 217,172,852 | C | T | 7 | 12 | maternal | 0 | 0 | ncRNA intronic |
| scaffold 1 | 219,008,542 | C | T | 9 | 14 | | 0 | 0 | intergenic |
| scaffold 1 | 220,218,852 | C | T | 9 | | | 0 | 0 | intronic |
| scaffold 1 | 229,516,801 | G | A | 7 | 13 | | 0 | 0 | intergenic |
| scaffold 1 | 241,925,043 | T | A | 10 | | | 0 | 0 | intergenic |
| scaffold 1 | 252,448,755 | G | A | 5 | | | 0 | 0 | intronic |
| scaffold 1 | 265,145,888 | G | A | 10 | | | 1 | 0 | intronic |
| scaffold 1 | 266,007,606 | G | C | 9 | | | 0 | 0 | intergenic |
| scaffold 1 | 270,609,098 | A | G | 10 | | | 0 | 0 | intergenic |
| scaffold 1 | 281,458,228 | C | T | 7 | 12 | | 0 | 1 | exonic |
| scaffold 1 | 281,822,529 | G | A | 6 | | | 0 | 0 | intergenic |
| scaffold 1 | 312,604,693 | C | T | 9 | 14 | | 0 | 1 | intronic |
| scaffold 2 | 1,865,246 | A | G | 6 | | | 0 | 0 | intergenic |
| scaffold 2 | 4,142,066 | G | C | 9 | 14 | | 1 | 0 | intronic |
| scaffold 2 | 5,142,496 | C | A | 11 | | | 0 | 0 | downstream |
| scaffold 2 | 7,708,293 | G | A | 9 | 14 | | 0 | 0 | intergenic |
| scaffold 2 | 8,288,259 | C | T | 9 | 14 | | 0 | 1 | intergenic |
| scaffold 2 | 14,303,251 | G | A | 8 | 12,13,14 | paternal | 1 | 0 | intronic |

| | | | | | | | | | |
|---|---|---|---|---|---|---|---|---|---|
| scaffold 2 | 14,608,975 | G | C | 10 | | | 0 | 0 | intergenic |
| scaffold 2 | 30,812,216 | A | G | 7 | 13 | paternal | 1 | 0 | intronic |
| scaffold 2 | 34,160,459 | T | G | 5 | | | 1 | 0 | intronic |
| scaffold 2 | 34,250,839 | C | T | 5 | | | 0 | 0 | intronic |
| scaffold 2 | 44,931,466 | G | A | 9 | | | 1 | 0 | ncRNA exonic |
| scaffold 2 | 54,599,969 | C | T | 8 | 12,13,14 | paternal | 0 | 1 | intergenic |
| scaffold 2 | 60,822,757 | G | A | 10 | | | 1 | 0 | intronic |
| scaffold 2 | 63,695,546 | G | A | 11 | | | 0 | 0 | intronic |
| scaffold 2 | 67,438,462 | A | G | 11 | | | 0 | 0 | intergenic |
| scaffold 2 | 91,579,546 | G | A | 10 | | | 0 | 1 | intronic |
| scaffold 2 | 93,530,505 | G | A | 6 | | | 0 | 0 | intronic |
| scaffold 2 | 105,283,328 | C | T | 9 | | | 0 | 0 | intronic |
| scaffold 2 | 125,877,124 | T | C | 9 | 14 | | 0 | 0 | intergenic |
| scaffold 2 | 134,591,113 | G | A | 9 | | | 0 | 0 | intronic |
| scaffold 2 | 134,592,401 | A | C | 9 | | | 0 | 0 | intronic |
| scaffold 2 | 142,178,571 | G | A | 11 | | | 0 | 0 | intronic |
| scaffold 2 | 154,564,808 | T | C | 11 | | | 0 | 0 | intronic |
| scaffold 2 | 160,014,264 | G | A | 7 | 13 | paternal | 0 | 1 | intronic |
| scaffold 2 | 171,382,931 | G | A | 10 | | | 1 | 1 | intergenic |
| scaffold 2 | 179,770,975 | G | A | 11 | | | 0 | 0 | intronic |
| scaffold 2 | 190,103,501 | G | A | 10 | | | 0 | 0 | intronic |
| scaffold 2 | 193,223,025 | G | C | 9 | 14 | | 0 | 0 | intergenic |
| scaffold 2 | 214,273,209 | T | G | 9 | 14 | | 0 | 0 | intronic |
| scaffold 2 | 218,720,566 | G | A | 11 | | | 1 | 0 | intergenic |
| scaffold 2 | 220,142,203 | T | C | 8 | 13,14 | paternal | 0 | 0 | ncRNA intronic |
| scaffold 2 | 231,186,418 | A | G | 8 | 12 | maternal | 0 | 0 | ncRNA intronic |
| scaffold 2 | 244,553,961 | G | T | 9 | | | 0 | 0 | upstream |
| scaffold 2 | 246,010,259 | G | A | 11 | | | 0 | 1 | intronic |
| scaffold 2 | 248,481,780 | C | T | 11 | | | 0 | 0 | intergenic |
| scaffold 2 | 250,162,294 | C | T | 6 | | | 1 | 0 | intronic |
| scaffold 2 | 254,341,502 | C | T | 11 | | | 0 | 0 | intronic |
| scaffold 2 | 273,627,907 | C | T | 9 | | | 0 | 1 | intergenic |
| scaffold 2 | 275,305,453 | G | T | 11 | | | 1 | 0 | intronic |
| scaffold 2 | 278,289,624 | G | A | 11 | | | 0 | 0 | ncRNA intronic |
| scaffold 2 | 281,345,131 | G | A | 11 | | | 0 | 1 | intergenic |
| scaffold 2 | 285,794,831 | G | A | 9 | | | 0 | 1 | intergenic |
| scaffold 3 | 477,012 | C | A | 9 | 14 | | 1 | 0 | intergenic |
| scaffold 3 | 12,903,626 | C | A | 7 | 12 | paternal | 0 | 0 | intergenic |
| scaffold 3 | 24,162,721 | T | G | 10 | | | 1 | 0 | intergenic |
| scaffold 3 | 32,519,996 | T | C | 5 | | | 0 | 0 | intergenic |
| scaffold 3 | 42,548,383 | C | T | 8 | 12 | paternal | 1 | 1 | intronic |
| scaffold 3 | 43,398,989 | T | C | 10 | | | 0 | 0 | intergenic |
| scaffold 3 | 52,418,893 | T | C | 9 | 14 | | 0 | 0 | intergenic |
| scaffold 3 | 61,464,118 | G | A | 11 | | | 0 | 0 | intergenic |
| scaffold 3 | 62,574,023 | G | A | 6 | | | 0 | 1 | ncRNA intronic |
| scaffold 3 | 66,343,255 | T | C | 9 | | | 0 | 0 | intergenic |
| scaffold 3 | 78,388,949 | T | C | 10 | | | 0 | 0 | intronic |
| scaffold 3 | 85,695,538 | A | G | 8 | 12 | paternal | 0 | 0 | intronic |
| scaffold 3 | 90,488,921 | G | A | 9 | 14 | | 0 | 0 | intergenic |
| scaffold 3 | 93,675,285 | C | T | 10 | | | 1 | 0 | intergenic |
| scaffold 3 | 96,623,599 | C | T | 11 | | | 0 | 0 | intergenic |
| scaffold 3 | 117,285,602 | C | T | 9 | | | 1 | 1 | ncRNA intronic |
| scaffold 3 | 141,612,598 | G | A | 10 | | | 1 | 0 | intergenic |
| scaffold 3 | 143,355,183 | T | C | 10 | | | 1 | 0 | intergenic |
| scaffold 3 | 148,740,638 | T | C | 7 | 13 | maternal | 1 | 0 | intergenic |
| scaffold 3 | 149,022,487 | T | C | 11 | | | 0 | 0 | intergenic |
| scaffold 3 | 160,874,583 | T | C | 10 | | | 1 | 0 | intergenic |
| scaffold 3 | 161,449,352 | G | A | 7 | | | 0 | 1 | intergenic |
| scaffold 3 | 168,733,956 | A | G | 9 | 14 | | 1 | 0 | intergenic |

| | | | | | | | | | |
|---|---|---|---|---|---|---|---|---|---|
| scaffold 3 | 175,256,215 | C | G | 9 | 14 | | 0 | 0 | intergenic |
| scaffold 3 | 175,256,776 | C | T | 9 | 14 | | 0 | 1 | intergenic |
| scaffold 3 | 175,256,940 | G | A | 9 | 14 | | 1 | 0 | intergenic |
| scaffold 3 | 199,324,638 | C | T | 7 | | | 0 | 1 | ncRNA intronic |
| scaffold 3 | 201,241,484 | C | T | 11 | | | 0 | 0 | intronic |
| scaffold 3 | 202,473,494 | C | T | 9 | 14 | | 0 | 0 | intergenic |
| scaffold 3 | 207,170,772 | G | A | 9 | 14 | | 1 | 0 | intronic |
| scaffold 3 | 232,600,418 | C | T | 10 | | | 1 | 0 | intergenic |
| scaffold 3 | 234,107,113 | G | A | 10 | | | 0 | 0 | intronic |
| scaffold 3 | 245,457,240 | A | T | 5 | | | 0 | 0 | intergenic |
| scaffold 3 | 248,530,341 | C | T | 10 | | | 0 | 0 | intergenic |
| scaffold 3 | 250,714,461 | C | A | 9 | 14 | | 0 | 0 | intronic |
| scaffold 3 | 257,539,258 | T | G | 10 | | | 1 | 0 | intergenic |
| scaffold 3 | 260,438,641 | C | G | 10 | | | 0 | 0 | intronic |
| scaffold 4 | 4,192,816 | G | A | 5 | | | 1 | 1 | ncRNA intronic |
| scaffold 4 | 10,942,451 | T | C | 11 | | | 0 | 0 | intergenic |
| scaffold 4 | 19,882,375 | G | A | 11 | | | 0 | 0 | intergenic |
| scaffold 4 | 20,711,847 | C | T | 7 | 12,13 | paternal | 1 | 0 | intronic |
| scaffold 4 | 22,984,960 | A | G | 7 | 12,13 | paternal | 1 | 0 | intronic |
| scaffold 4 | 24,260,188 | C | T | 5 | | | 0 | 0 | intergenic |
| scaffold 4 | 26,941,045 | A | T | 9 | 14 | | 0 | 0 | intronic |
| scaffold 4 | 27,975,708 | C | T | 11 | | | 1 | 1 | intronic |
| scaffold 4 | 28,627,101 | C | T | 6 | | | 1 | 0 | intronic |
| scaffold 4 | 40,316,647 | T | G | 5 | | | 0 | 0 | intronic |
| scaffold 4 | 56,113,788 | C | G | 7 | 12,13 | paternal | 1 | 0 | intergenic |
| scaffold 4 | 73,767,575 | C | T | 7 | 12,13 | paternal | 1 | 0 | intronic |
| scaffold 4 | 83,689,499 | G | A | 10 | | | 1 | 0 | intergenic |
| scaffold 4 | 84,507,725 | T | G | 8 | 13,14 | maternal | 0 | 0 | intergenic |
| scaffold 4 | 89,126,688 | A | G | 11 | | | 1 | 0 | intergenic |
| scaffold 4 | 94,466,741 | C | T | 9 | 14 | | 0 | 0 | ncRNA intronic |
| scaffold 4 | 102,874,903 | A | C | 8 | 12 | paternal | 1 | 0 | ncRNA intronic |
| scaffold 4 | 112,894,534 | A | T | 9 | 14 | | 0 | 0 | intergenic |
| scaffold 4 | 113,119,213 | G | A | 8 | 12 | paternal | 0 | 0 | intergenic |
| scaffold 4 | 114,169,135 | G | A | 7 | 12,13 | | 0 | 0 | intronic |
| scaffold 4 | 115,093,271 | C | T | 5 | | | 0 | 0 | intronic |
| scaffold 4 | 137,233,600 | G | T | 6 | | | 1 | 0 | intergenic |
| scaffold 4 | 149,274,846 | A | G | 10 | | | 0 | 0 | intergenic |
| scaffold 4 | 155,468,886 | T | C | 9 | 14 | | 0 | 0 | ncRNA intronic |
| scaffold 4 | 156,772,407 | G | A | 5 | | | 0 | 1 | intronic |
| scaffold 4 | 157,446,185 | C | T | 10 | | | 1 | 0 | intergenic |
| scaffold 4 | 167,950,896 | T | A | 10 | | | 0 | 0 | UTR3 |
| scaffold 4 | 172,193,529 | G | A | 9 | 14 | | 0 | 0 | intronic |
| scaffold 4 | 180,633,547 | A | G | 7 | | | 1 | 0 | intergenic |
| scaffold 4 | 185,111,613 | G | A | 8 | | | 0 | 0 | intergenic |
| scaffold 4 | 191,157,805 | G | A | 9 | | | 0 | 1 | intronic |
| scaffold 4 | 191,329,469 | A | G | 10 | | | 1 | 0 | intronic |
| scaffold 4 | 194,273,261 | T | C | 10 | | | 0 | 0 | intergenic |
| scaffold 4 | 196,460,955 | T | A | 11 | | | 0 | 0 | intergenic |
| scaffold 4 | 198,689,245 | C | A | 9 | 14 | | 1 | 0 | intergenic |
| scaffold 4 | 198,732,991 | A | G | 10 | | | 0 | 0 | intergenic |
| scaffold 4 | 208,177,391 | A | G | 11 | | | 0 | 0 | intergenic |
| scaffold 4 | 213,802,032 | G | A | 9 | | | 0 | 0 | intergenic |
| scaffold 5 | 3,801,567 | G | A | 6 | | | 1 | 0 | intronic |
| scaffold 5 | 8,219,036 | T | C | 9 | | | 0 | 0 | intronic |
| scaffold 5 | 25,758,442 | C | T | 11 | | | 1 | 0 | intergenic |
| scaffold 5 | 26,120,875 | G | A | 6 | | | 0 | 1 | intronic |
| scaffold 5 | 35,585,107 | C | T | 9 | | | 1 | 1 | intergenic |
| scaffold 5 | 36,245,980 | G | A | 5 | | | 1 | 0 | intergenic |
| scaffold 5 | 53,600,800 | T | C | 7 | 13 | paternal | 0 | 0 | intergenic |

| | | | | | | | | | |
|---|---|---|---|---|---|---|---|---|---|
| scaffold 5 | 60,814,071 | A | G | 9 | 14 | | 1 | 0 | intergenic |
| scaffold 5 | 68,547,849 | A | G | 6 | | | 0 | 0 | intronic |
| scaffold 5 | 69,944,542 | T | A | 7 | 13 | paternal | 0 | 0 | intronic |
| scaffold 5 | 69,958,324 | T | A | 11 | | | 1 | 0 | intronic |
| scaffold 5 | 78,176,561 | C | T | 5 | | | 1 | 0 | intronic |
| scaffold 5 | 101,402,050 | C | G | 11 | | | 1 | 0 | intergenic |
| scaffold 5 | 112,064,169 | T | G | 9 | 14 | | 1 | 0 | intergenic |
| scaffold 5 | 116,186,489 | A | C | 5 | | | 0 | 0 | intronic |
| scaffold 5 | 117,084,450 | A | G | 11 | | | 1 | 0 | intergenic |
| scaffold 5 | 117,283,892 | T | C | 5 | | | 0 | 0 | intergenic |
| scaffold 5 | 127,622,924 | G | C | 10 | | | 0 | 0 | ncRNA intronic |
| scaffold 5 | 131,636,766 | C | T | 10 | | | 0 | 0 | exonic |
| scaffold 5 | 135,282,560 | C | T | 5 | | | 0 | 0 | intronic |
| scaffold 5 | 135,730,394 | G | A | 11 | | | 0 | 0 | intergenic |
| scaffold 5 | 146,034,809 | C | T | 10 | | | 1 | 1 | intergenic |
| scaffold 5 | 148,255,879 | T | C | 10 | | | 1 | 0 | intronic |
| scaffold 5 | 165,742,306 | T | C | 10 | | | 0 | 0 | intronic |
| scaffold 5 | 178,910,388 | T | A | 10 | | | 0 | 0 | intergenic |
| scaffold 5 | 181,580,493 | C | G | 9 | | | 1 | 0 | intergenic |
| scaffold 5 | 185,864,401 | C | T | 9 | 14 | | 1 | 0 | intronic |
| scaffold 5 | 187,869,366 | A | G | 8 | 12,14 | paternal | 0 | 0 | intergenic |
| scaffold 5 | 194,842,877 | G | A | 6,8 | | | 1 | 1 | intronic |
| scaffold 5 | 197,829,626 | G | A | 5 | | | 0 | 1 | intronic |
| scaffold 5 | 206,567,735 | G | T | 10 | | | 0 | 0 | intronic |
| scaffold 5 | 207,698,431 | G | A | 11 | | | 0 | 1 | intergenic |
| scaffold 6 | 19,709,982 | T | C | 9 | 14 | | 1 | 0 | intergenic |
| scaffold 6 | 22,788,684 | C | T | 6 | | | 0 | 0 | upstream |
| scaffold 6 | 23,766,882 | G | A | 11 | | | 0 | 0 | ncRNA exonic |
| scaffold 6 | 25,870,729 | C | T | 9 | 14 | | 0 | 0 | intergenic |
| scaffold 6 | 30,024,772 | G | A | 10 | | | 1 | 0 | intergenic |
| scaffold 6 | 37,732,979 | G | A | 7 | 13 | | 1 | 0 | intronic |
| scaffold 6 | 60,415,268 | T | C | 11 | | | 0 | 0 | intergenic |
| scaffold 6 | 66,053,686 | G | A | 10 | | | 1 | 1 | intergenic |
| scaffold 6 | 75,005,246 | C | T | 5 | | | 0 | 0 | intergenic |
| scaffold 6 | 82,984,459 | T | C | 11 | | | 0 | 0 | intergenic |
| scaffold 6 | 93,859,481 | T | C | 5 | | | 0 | 0 | ncRNA intronic |
| scaffold 6 | 107,315,614 | G | A | 5 | | | 1 | 0 | intronic |
| scaffold 6 | 119,404,902 | G | A | 10 | | | 1 | 1 | intergenic |
| scaffold 6 | 119,452,340 | C | T | 5 | | | 0 | 0 | intergenic |
| scaffold 6 | 120,133,712 | C | G | 10 | | | 0 | 0 | intergenic |
| scaffold 6 | 126,977,559 | T | C | 8 | | | 1 | 0 | ncRNA exonic |
| scaffold 6 | 127,023,149 | C | A | 10 | | | 1 | 0 | ncRNA intronic |
| scaffold 6 | 134,734,524 | G | T | 8 | 12,13,14 | maternal | 1 | 0 | intergenic |
| scaffold 6 | 137,337,573 | G | A | 5 | | | 0 | 1 | exonic |
| scaffold 6 | 145,471,715 | G | A | 8 | | | 0 | 0 | intergenic |
| scaffold 6 | 146,916,039 | C | A | 9 | | | 0 | 0 | intergenic |
| scaffold 6 | 157,797,328 | A | G | 9 | 14 | | 1 | 0 | intronic |
| scaffold 6 | 158,266,570 | G | A | 11 | | | 0 | 0 | intergenic |
| scaffold 6 | 162,882,463 | G | A | 10 | | | 0 | 1 | ncRNA intronic |
| scaffold 6 | 182,730,862 | T | C | 6 | | | 0 | 0 | ncRNA intronic |
| scaffold 6 | 184,254,493 | T | C | 10 | | | 1 | 0 | intergenic |
| scaffold 6 | 192,239,354 | C | T | 8 | 13 | paternal | 1 | 0 | intergenic |
| scaffold 6 | 194,199,182 | G | A | 10 | | | 0 | 1 | intergenic |
| scaffold 6 | 197,268,148 | G | A | 11 | | | 0 | 0 | ncRNA exonic |
| scaffold 6 | 198,713,613 | C | T | 9 | 14 | | 0 | 1 | intergenic |
| scaffold 7 | 18,021,451 | C | T | 11 | | | 0 | 1 | intronic |
| scaffold 7 | 20,261,654 | G | A | 10 | | | 0 | 0 | intergenic |
| scaffold 7 | 47,353,130 | G | A | 10 | | | 0 | 1 | exonic |
| scaffold 7 | 48,918,875 | T | C | 10 | | | 1 | 0 | ncRNA intronic |

| | | | | | | | | | |
|---|---|---|---|---|---|---|---|---|---|
| scaffold 7 | 53,085,677 | T | C | 9 | 14 | | 0 | 0 | ncRNA intronic |
| scaffold 7 | 54,991,651 | T | G | 11 | | | 1 | 0 | intronic |
| scaffold 7 | 65,197,835 | G | A | 10 | | | 0 | 0 | intronic |
| scaffold 7 | 74,876,778 | C | G | 11 | | | 0 | 0 | intronic |
| scaffold 7 | 83,798,482 | A | C | 9 | | | 0 | 0 | UTR3 |
| scaffold 7 | 85,202,968 | A | G | 9 | | | 0 | 0 | intronic |
| scaffold 7 | 91,485,433 | G | T | 11 | | | 0 | 0 | ncRNA intronic |
| scaffold 7 | 96,649,799 | A | G | 10 | | | 0 | 0 | intergenic |
| scaffold 7 | 97,115,933 | T | A | 7 | | | 0 | 0 | ncRNA intronic |
| scaffold 7 | 98,906,215 | G | C | 10 | | | 0 | 0 | intronic |
| scaffold 7 | 102,052,828 | T | C | 9 | 14 | | 0 | 0 | intergenic |
| scaffold 7 | 106,843,062 | A | G | 10 | | | 0 | 0 | ncRNA intronic |
| scaffold 7 | 109,907,860 | C | T | 8 | 12,14 | paternal | 0 | 0 | intergenic |
| scaffold 7 | 110,813,748 | G | A | 11 | | | 1 | 0 | intergenic |
| scaffold 7 | 113,953,090 | G | A | 10 | | | 1 | 0 | ncRNA intronic |
| scaffold 7 | 136,189,002 | T | C | 5 | | | 1 | 0 | ncRNA intronic |
| scaffold 7 | 138,454,011 | T | A | 8 | 12,14 | paternal | 1 | 0 | ncRNA intronic |
| scaffold 7 | 160,364,697 | G | A | 7 | | | 1 | 0 | intronic |
| scaffold 7 | 170,616,616 | G | A | 7 | | | 0 | 1 | ncRNA intronic |
| scaffold 7 | 173,544,704 | G | A | 11 | | | 1 | 1 | ncRNA intronic |
| scaffold 7 | 173,815,640 | A | G | 10 | | | 1 | 0 | intronic |
| scaffold 7 | 176,907,736 | C | T | 6 | | | 0 | 1 | intergenic |
| scaffold 7 | 188,829,124 | C | T | 9 | 14 | | 0 | 0 | intronic |
| scaffold 7 | 190,816,781 | G | A | 10 | | | 0 | 0 | intergenic |
| scaffold 8 | 2,591,834 | C | T | 9 | | | 0 | 0 | intergenic |
| scaffold 8 | 7,763,347 | G | A | 11 | | | 0 | 0 | intergenic |
| scaffold 8 | 8,211,116 | G | A | 5 | | | 0 | 1 | intergenic |
| scaffold 8 | 15,661,066 | A | G | 10 | | | 0 | 0 | intronic |
| scaffold 8 | 17,619,032 | C | A | 8 | 12,14 | paternal | 1 | 0 | intergenic |
| scaffold 8 | 28,810,684 | G | C | 5 | | | 0 | 0 | intronic |
| scaffold 8 | 46,687,622 | C | A | 10 | | | 0 | 0 | intronic |
| scaffold 8 | 49,661,418 | A | T | 10 | | | 0 | 0 | intronic |
| scaffold 8 | 57,434,258 | C | T | 11 | | | 0 | 1 | intergenic |
| scaffold 8 | 61,657,702 | C | T | 9 | | | 0 | 1 | intronic |
| scaffold 8 | 64,381,619 | G | A | 7 | 13 | maternal | 1 | 1 | ncRNA intronic |
| scaffold 8 | 79,462,092 | C | T | 5 | | | 0 | 1 | intronic |
| scaffold 8 | 85,608,633 | A | G | 10 | | | 1 | 0 | intergenic |
| scaffold 8 | 98,069,734 | G | A | 9 | | | 0 | 1 | ncRNA intronic |
| scaffold 8 | 106,401,209 | C | T | 10 | | | 0 | 1 | intronic |
| scaffold 8 | 108,865,160 | T | A | 10 | | | 0 | 0 | intergenic |
| scaffold 8 | 118,172,862 | T | G | 8 | 12,13 | paternal | 0 | 0 | upstream |
| scaffold 8 | 120,465,541 | C | T | 11 | | | 0 | 0 | intergenic |
| scaffold 8 | 154,181,996 | T | A | 10 | | | 0 | 0 | intergenic |
| scaffold 8 | 159,194,818 | C | T | 11 | | | 1 | 0 | intergenic |
| scaffold 8 | 162,669,441 | G | A | 9 | 14 | | 0 | 0 | intronic |
| scaffold 10 | 24,308,126 | G | A | 11 | | | 1 | 1 | intronic |
| scaffold 10 | 39,269,490 | G | A | 9 | 14 | | 0 | 0 | intergenic |
| scaffold 10 | 41,548,089 | G | A | 11 | | | 0 | 1 | intronic |
| scaffold 10 | 49,038,308 | A | G | 7 | | | 0 | 0 | UTR5 |
| scaffold 10 | 54,051,664 | T | C | 11 | | | 0 | 0 | ncRNA intronic |
| scaffold 10 | 75,655,410 | G | A | 9 | | | 0 | 0 | intergenic |
| scaffold 10 | 83,159,359 | C | T | 7 | 12,13 | paternal | 0 | 1 | intergenic |
| scaffold 10 | 98,841,732 | T | C | 10 | | | 0 | 0 | ncRNA intronic |
| scaffold 10 | 102,037,580 | C | A | 9 | 14 | | 0 | 0 | ncRNA intronic |
| scaffold 10 | 107,444,269 | G | A | 9 | 14 | | 1 | 0 | intronic |
| scaffold 10 | 114,150,198 | C | T | 9 | | | 0 | 0 | upstream |
| scaffold 11 | 233,973 | T | C | 10 | | | 1 | 0 | intronic |
| scaffold 11 | 20,715,958 | A | G | 10 | | | 0 | 0 | intronic |
| scaffold 11 | 31,093,279 | T | C | 11 | | | 0 | 0 | intronic |

| | | | | | | | | | |
|---|---|---|---|---|---|---|---|---|---|
| scaffold 11 | 34,632,852 | C | T | 5 | | | 0 | 0 | intronic |
| scaffold 11 | 40,766,617 | G | A | 8 | 13,14 | maternal | 1 | 1 | intergenic |
| scaffold 11 | 41,948,855 | C | T | 10 | | | 0 | 0 | downstream |
| scaffold 11 | 65,422,259 | T | G | 9 | 14 | | 0 | 0 | intronic |
| scaffold 11 | 67,455,235 | G | A | 10 | | | 0 | 0 | intergenic |
| scaffold 11 | 67,570,950 | C | T | 8 | | | 0 | 0 | intronic |
| scaffold 11 | 67,601,405 | A | G | 9 | | | 0 | 0 | intergenic |
| scaffold 11 | 81,668,502 | C | T | 11 | | | 0 | 0 | intronic |
| scaffold 11 | 82,887,976 | G | A | 9 | | | 0 | 0 | intronic |
| scaffold 11 | 84,030,543 | A | G | 10 | | | 1 | 0 | intergenic |
| scaffold 11 | 84,828,978 | C | T | 11 | | | 0 | 1 | intergenic |
| scaffold 11 | 85,969,795 | T | C | 10 | | | 0 | 0 | intronic |
| scaffold 11 | 87,560,176 | C | T | 9 | | | 0 | 0 | intronic |
| scaffold 11 | 93,822,210 | A | G | 11 | | | 1 | 0 | intronic |
| scaffold 11 | 94,458,460 | C | T | 7 | 12 | maternal | 1 | 0 | intergenic |
| scaffold 12 | 6,865,288 | G | A | 10 | | | 0 | 0 | intergenic |
| scaffold 12 | 7,299,069 | A | T | 10 | | | 0 | 0 | intergenic |
| scaffold 12 | 33,894,664 | G | A | 7 | 12,13 | paternal | 0 | 0 | intronic |
| scaffold 12 | 42,421,551 | A | T | 9 | | | 0 | 0 | intronic |
| scaffold 12 | 60,525,190 | T | G | 10 | | | 1 | 0 | intergenic |
| scaffold 12 | 60,559,798 | C | T | 10 | | | 1 | 0 | intergenic |
| scaffold 13 | 5,473,072 | C | T | 10 | | | 1 | 0 | ncRNA intronic |
| scaffold 13 | 17,138,170 | C | T | 11 | | | 0 | 0 | intergenic |
| scaffold 13 | 43,605,059 | T | C | 10 | | | 1 | 0 | intergenic |
| scaffold 13 | 44,628,884 | T | C | 8 | | | 1 | 0 | intronic |
| scaffold 13 | 61,045,460 | G | C | 10 | | | 0 | 0 | intergenic |
| scaffold 13 | 62,444,270 | G | A | 9 | | | 1 | 0 | intergenic |
| scaffold 14 | 9,751,324 | G | T | 9 | | | 0 | 0 | intergenic |
| scaffold 14 | 12,961,134 | T | C | 7 | 13 | paternal | 0 | 0 | intronic |
| scaffold 15 | 7,020,564 | A | T | 9 | | | 1 | 0 | ncRNA intronic |
| scaffold 15 | 11,703,775 | A | G | 11 | | | 0 | 0 | intergenic |
| scaffold 15 | 14,904,944 | A | T | 9 | 14 | | 1 | 0 | intronic |
| scaffold 15 | 18,873,275 | C | T | 11 | | | 0 | 0 | intronic |
| scaffold 15 | 19,230,056 | G | A | 10 | | | 0 | 1 | intronic |
| scaffold 15 | 23,088,485 | C | T | 9 | | | 0 | 0 | intronic |
| scaffold 15 | 26,916,571 | C | T | 9 | | | 0 | 1 | intergenic |

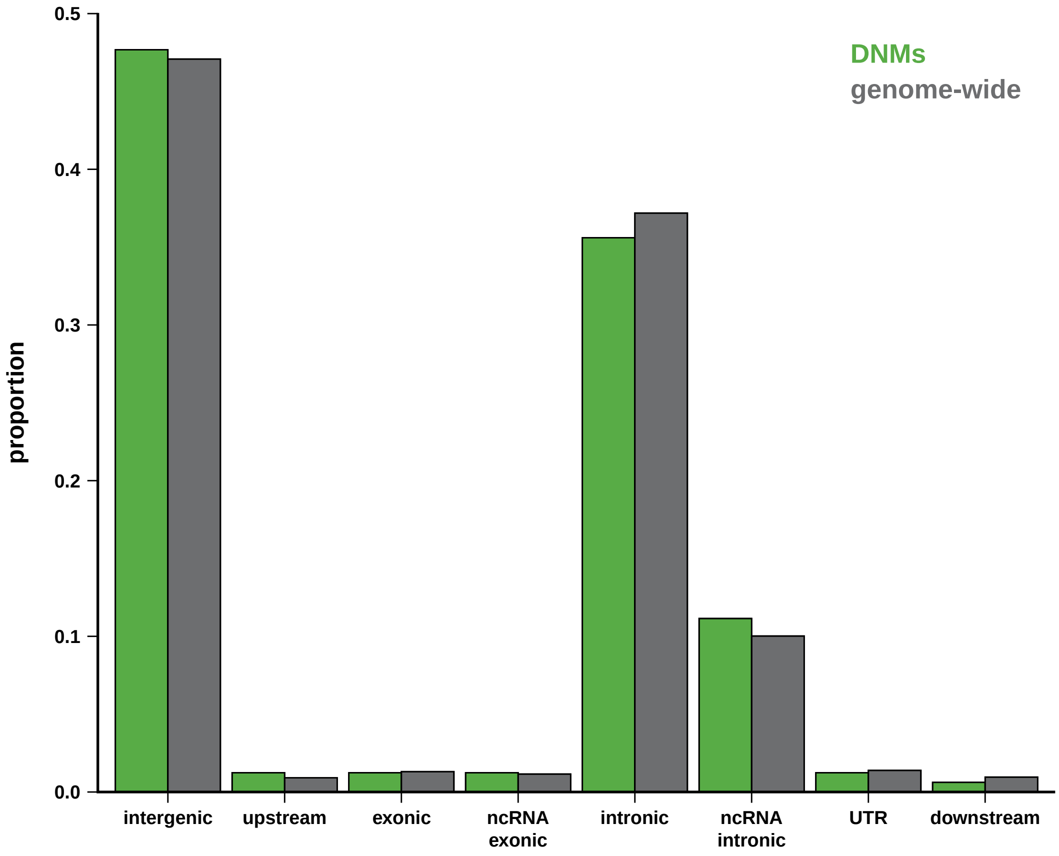


**Supplementary Figure 1**. Distribution of *de novo* mutations (DNMs; shown in green) as well as sites accessible to the study (gray) by genomic region (*i.e.*, intergenic, upstream, exonic, exonic non-coding RNA [ncRNA], intronic, intronic ncRNA, 3' and 5' UTR, and downstream). Genome annotations were obtained from the the aye-aye genome assembly (Versoza and Pfeifer 2024).

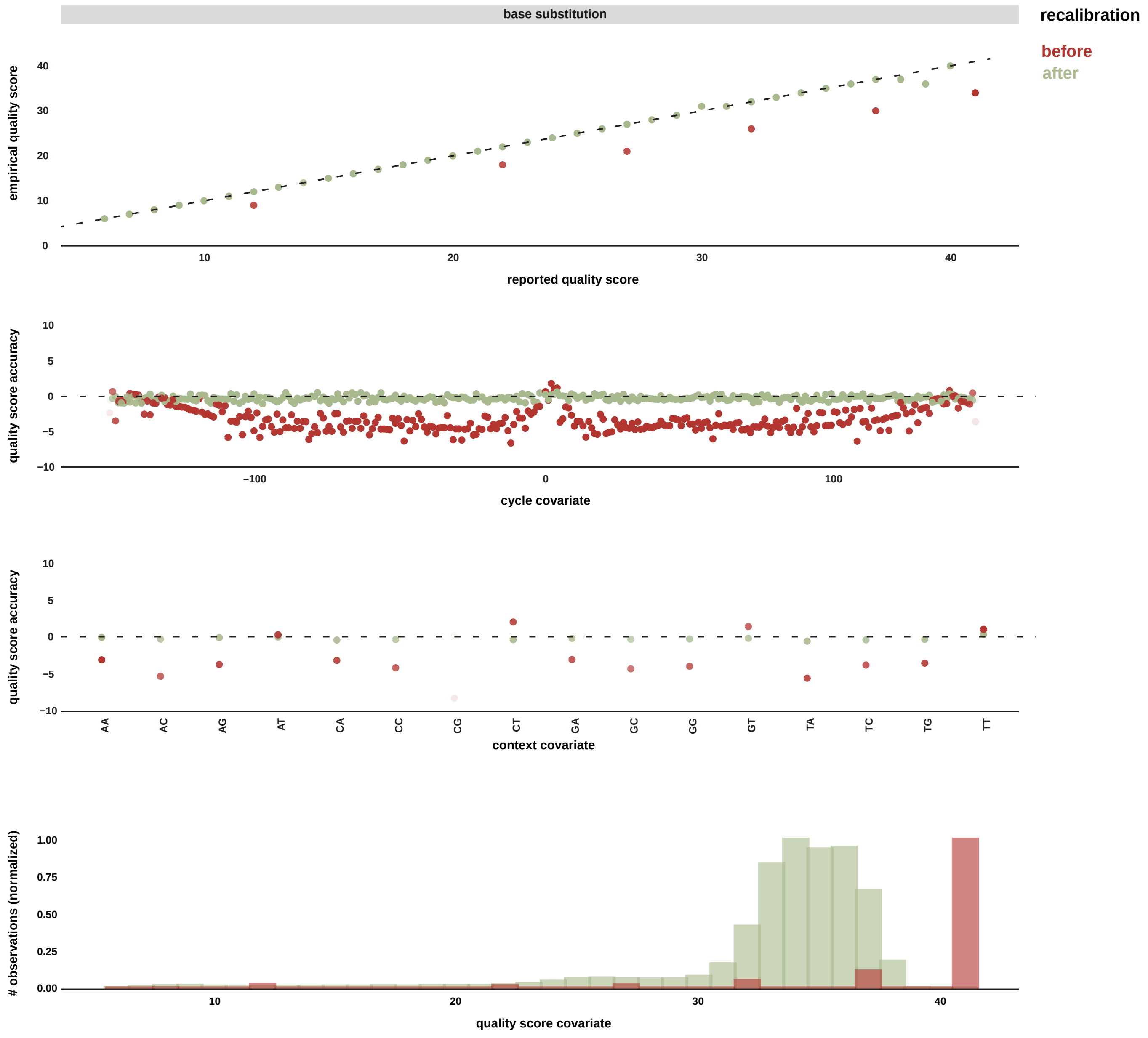


**Supplementary Figure 2**. Quality scores before (shown in red) and after (green) Base Quality Score Recalibration (BQSR).

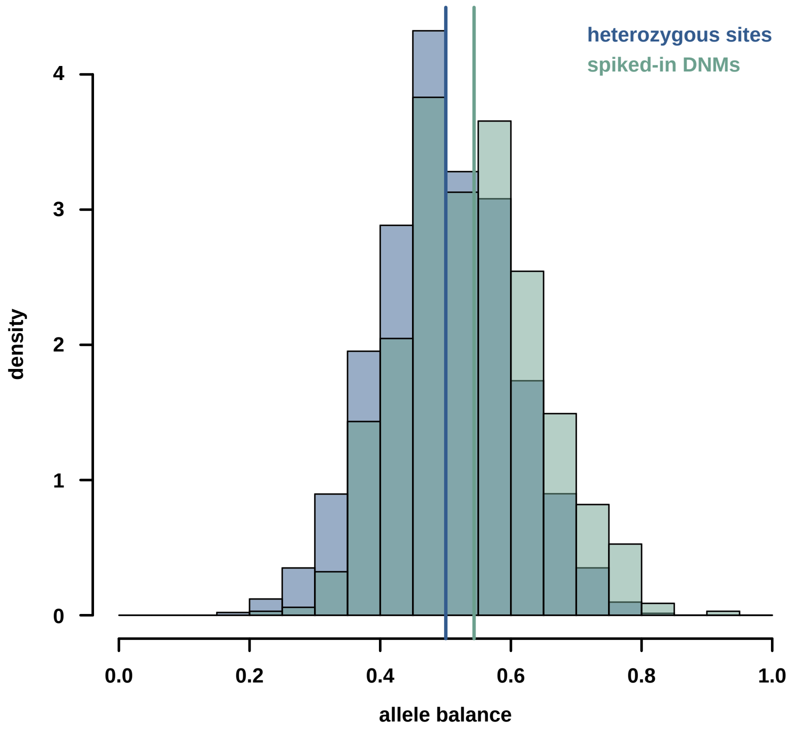


**Supplementary Figure 3**. Distribution of the allele balance (i.e., the ratio of reads carrying the alternative vs reference alleles) at genuine heterozygous sites (shown in blue) as well as at the simulated (spiked-in) *de novo* mutations (DNMs; shown in teal).
